# Multiple phase-variable mechanisms, including capsular polysaccharides, modify bacteriophage susceptibility in *Bacteroides thetaiotaomicron*

**DOI:** 10.1101/521070

**Authors:** Nathan T. Porter, Andrew J. Hryckowian, Bryan D. Merrill, Jaime J. Fuentes, Jackson O. Gardner, Robert W. P. Glowacki, Shaleni Singh, Ryan D. Crawford, Evan S. Snitkin, Justin L. Sonnenburg, Eric C. Martens

**Author notes:** These authors contributed equally to this work (Nathan T. Porter, Andrew J. Hryckowian).

## Abstract

A variety of cell surface structures, including capsular polysaccharides (CPS), dictate interactions between bacteria and their environment including their viruses (bacteriophages). Members of the prominent human gut Bacteroidetes characteristically produce several phase-variable CPS, but their contributions to bacteriophage interactions are unknown. We used engineered strains of the human symbiont *Bacteroides thetaiotaomicron*, which differ only in the CPS they express, to isolate bacteriophages from two locations in the United States. Testing each of 71 bacteriophages against a panel of strains that express wild-type phase-variable CPS, one of eight different single CPS, or no CPS at all, revealed that each phage infects only a subset of otherwise isogenic strains. Deletion of infection-permissive CPS from *B. thetaiotaomicron* was sufficient to abolish infection for several individual bacteriophages, while infection of wild-type *B. thetaiotaomicron* with either of two different bacteriophages rapidly selected for expression of non-permissive CPS. Surprisingly, acapsular *B. thetaiotaomicron* also escapes complete killing by these bacteriophages, but surviving bacteria exhibit increased expression of 8 distinct phase-variable lipoproteins. When constitutively expressed, one of these lipoproteins promotes resistance to multiple bacteriophages. Finally, both wild-type and acapsular *B. thetaiotaomicron* were able to separately co-exist with one bacteriophage for over two months in the mouse gut, suggesting that phase-variation promotes resistance but also generates sufficient numbers of susceptible revertants to allow bacteriophage persistence. Our results reveal important roles for *Bacteroides* CPS and other cell surface structures that allow these bacteria to persist despite bacteriophage predation and hold important implications for using bacteriophages therapeutically to target gut symbionts.

## Introduction

The community of cellular microorganisms in the human intestinal tract is dominated by a diverse population of bacteria, with hundreds of different species typically coexisting within an individual^1, 2^. Frequent diet changes, host immune responses and bacteriophage infections are among the many causes of intermittent perturbations to individual bacterial taxa. However, the microbial communities within individuals generally remain stable over long time periods^3^, suggesting that bacteria have evolved strategies to survive these perturbations. One mechanism that may promote bacterial resilience is the ability of individual strains to produce multiple capsular polysaccharides (CPS), cell surface components that have been diversified in the genomes of gut-dwelling Bacteroidetes and several other phyla^4, 5^. While previous work showed that CPS from *Bacteroides* and members of other phyla play roles in evading or modulating host immunity^6–10^, the diversity of CPS synthesis loci in gut bacteria suggests that they could fill other roles^5, 8, 11, 12^.

The phylum Bacteroidetes—within which members of the genus *Bacteroides* are typically the most abundant Gram-negative gut symbionts in industrialized human populations^2, 13^—provides excellent models to study persistence and competition mechanisms, including the roles and diversity of CPS. For example, the type strains of the well-studied species *Bacteroides thetaiotaomicron* and *Bacteroides fragilis* each encode 8 different CPS^14, 15^ and there is broad genetic diversity of *cps* loci among different strains within these species (e.g., 47 different *cps* biosynthetic loci were identified in just 14 strains of *B. thetaiotaomicron*)^8^. In *Bacteroides*, CPS structures appear to surround the entire bacterial cell^16, 17^ and the *cps* biosynthetic loci that encode these surface coatings are often under the control of phase variable promoters^8, 15, 18^. In conjunction with other regulatory mechanisms, phase variable CPS expression generates phenotypic heterogeneity within an otherwise isogenic population that may facilitate survival in the face of diverse disturbances^8, 15, 19, 20^.

Bacterial viruses or bacteriophages (herein, phages), like the bacteria on which they prey, vary greatly across individual gut microbiomes and are even responsive to host dietary changes and disease states^21–25^. Compared to gut bacteria, far less is understood about the phages of the gut microbiome, especially the mechanisms governing phage-bacteria interactions. Specifically, while phages that target several species of *Bacteroides* have been shown to exhibit species- or strain-specificity^26–29^, little is known about the molecular interactions that drive bacterial susceptibility^30^ or the mechanisms by which these bacteria persist despite an abundance of phages in the gut. Given the observations that *Bacteroides* CPS are extremely variable, even within members of a single species^8, 11^, and employ complex regulatory mechanisms that diversify expression in members of a population^20^, CPS are ideal candidates for modulating *Bacteroides*-phage interactions.

Here, we tested the hypothesis that CPS mediate *Bacteroides*-phage interactions. We employed a panel of engineered strains of the model symbiont *Bacteroides thetaiotaomicron* that each constitutively expresses a different single CPS or none at all. While our results clearly support the conclusion that individual CPS can either block or be required for phage infection, they also reveal that *B. thetaiotaomicron* possesses additional phage-evasion strategies that function in addition to CPS. For two different phages tested, CPS-independent survival involves increased expression of phase-variable surface lipoproteins and altered expression of nutrient receptors by the surviving bacteria. These phase-variable surface proteins may also encode resistance mechanisms, an idea that is supported by increased resistance to several phages when one of these lipoproteins is constitutively expressed experimentally. Our results provide a mechanistic glimpse into the intricacy of bacterial-phage interactions that exist in the human gut and provide a foundation for future work to leverage these interactions and to directly manipulate the gut microbiome.

## Results

### Bacteriophages infect *B. thetaiotaomicron* in a CPS-dependent fashion

The genomes of human gut Bacteroidetes frequently encode multiple CPS^5^, a phenomenon that we explored in a phylogenetic context in currently available genomes of 53 different human gut species. The type strains of all but the *Prevotella* species searched encoded between 2-13 CPS (mean = 4), suggesting that the ability to produce multiple CPS is typical and some lineages have undergone substantial expansion of these surface structures (**Figure S1A**). To test the hypothesis that *Bacteroides* CPS mediate interactions with phages, we isolated phages that infect *B. thetaiotaomicron* VPI-5482 (ATCC 29148). To maximize our chances of collecting phages that differ in their interactions with CPS, we used the wild-type strain that expresses 8 different CPS, which are each encoded by a different multi-gene *cps* locus and in some cases are driven by phase-variable promoters (**Figure S1B**)^14^, along with a panel of engineered strains with reduced CPS expression. The latter included 8 single CPS-expressing strains (designated “cps1” through “cps8”)^8^ and an acapsular strain in which all eight *cps* loci were deleted (Δ*cps*)^31^ as independent hosts for phage isolation. Primary sewage effluent from two cities within the United States (Ann Arbor, Michigan and San Jose, California; separated by approximately 3,300 kilometers) was used as the phage source (for further details on phage isolation, see *Methods* and **Table S1**). All phages were plaque purified at least 3 times and high titer lysates generated for each of the 71 phages. Plaque morphologies varied among the individual phages, ranging in size from <1 mm to >3 mm and in opacity from very turbid to clear (**Figure S2**).

To determine if phages isolated on individual *B. thetaiotaomicron* hosts are restricted by the particular CPS they express, we systematically tested each phage against each of the 10 host strains (n=3). Hierarchical clustering of the host infection profiles revealed a cladogram with 3 main branches that each encompasses phages from both collection sites, although substantial variation in host tropism exists for phages within each branch (**Figure 1**). Furthermore, individual phages within each branch displayed a range of plaque morphologies (**Figures 1, S2**), suggesting additional diversity in the collection that is not captured by this assay. Finally, host range assays were robust when performed by different experimenters at different research sites (**Figure S3**).

**Figure 1.**
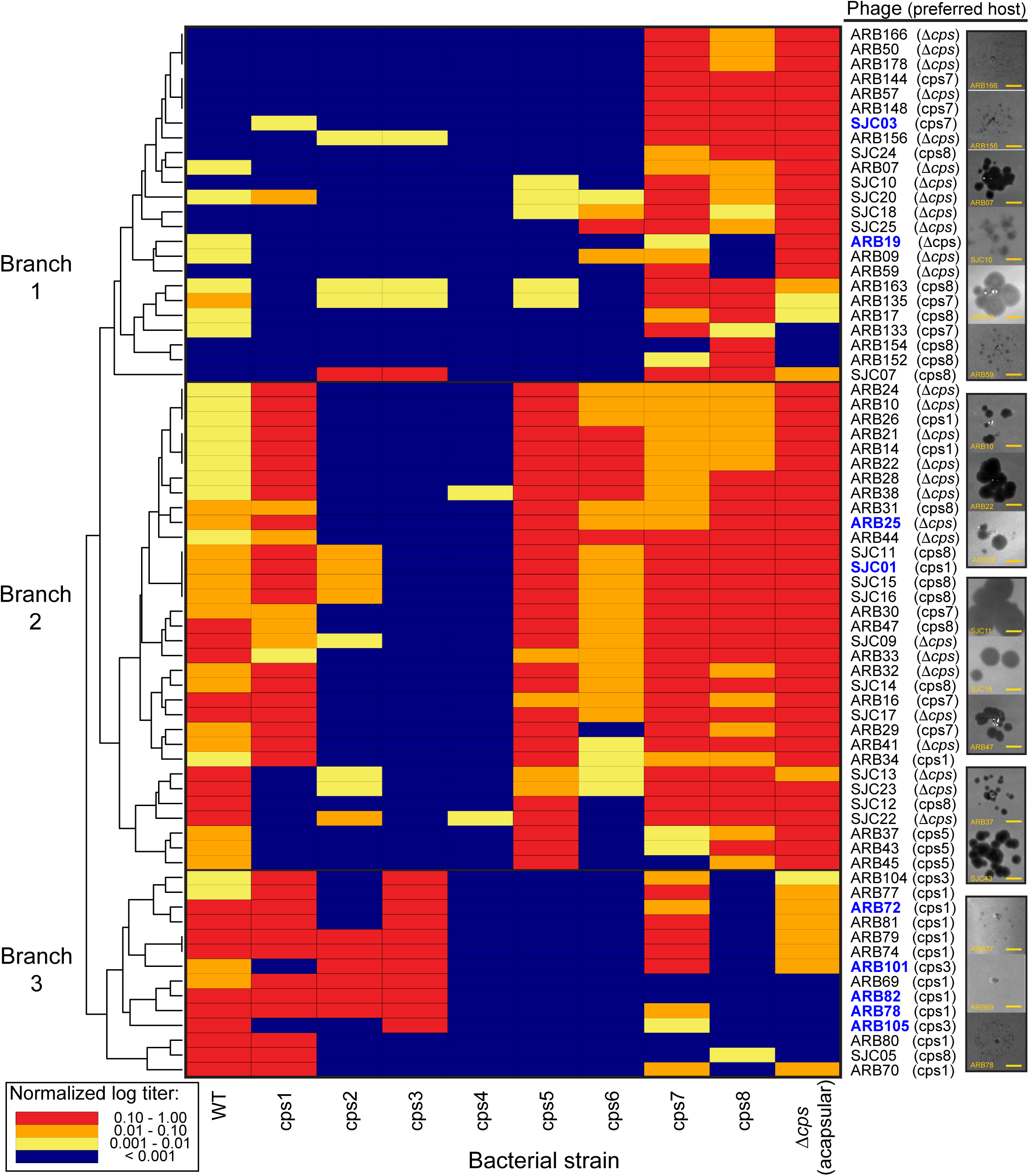
Host range of *B. thetaiotaomicron* phages on strains expressing different CPS. Seventy-one bacteriophages were isolated and purified on the wild-type, Δ*cps* (acapsular), or the 8 single CPS-expressing *B. thetaiotaomicron* strains. High titer phage stocks were prepared on their “preferred host strain”, which was the strain that yielded the highest titer of phages in a pre-screen of phage host range and is listed next to each phage. Phages were then tested in a quantitative host range assay. Phage titers were calculated for each bacterial host and normalized to the preferred host strain for each replicate, 3 replicates averaged for each assay and the results clustered based on plaquing efficiencies (see *Methods*). Images at the far right of the figure illustrate the range of plaque morphologies of select phages from the collection (see **Figure S2** for images of plaques for all phages). Several phages that are the subjects of additional follow up studies are highlighted in blue text. Scale bar = 2mm.

Phages in Branch 1 generally exhibited robust infection of the acapsular strain, although 3 of these phages did not form plaques on this host. Furthermore, phages in Branch 1 generally exhibited robust infection on strains expressing CPS7 or CPS8 alone, although a separate subset of 3 phages did not form plaques on the CPS8 expressing strain. Some Branch 1 phages also displayed less efficient infection of other strains with the exception of cps4, which was not infected by any phages in this group. Interestingly, ARB154 exclusively infected cps8, an uncommon CPS among *B. thetaiotaomicron* strains that appears to be contained in a mobile element^8^. Phages in Branch 2 generally exhibited robust infection of all strains except cps2, cps3 and cps4. However, subsets of this group were unable to infect cps1 or cps6. Finally, Branch 3 tended to exhibit strong infection of wild-type, *cps*1, cps2, and cps3, with some variations. Some Branch 3 phages also exhibited the ability to infect the cps7 and acapsular strains but were the only branch that poorly infected cps8. Most notably, a subset of phages on Branches 1 and 3 failed to infect the acapsular strain, suggesting that they require the presence of certain CPS for infection. Taken together, the observed variations in phage infectivity provide support for our hypothesis that these surface structures are important mediators of *B. thetaiotaomicron*-phage interactions.

### Elimination of specific CPS subsets alters bacterial susceptibility to phages

The differences in host infectivity described above suggest that there are distinct mechanisms of phage adsorption to the bacterial surface, some of which are influenced by CPS. Several phages robustly infect the acapsular strain, indicating that a capsule-independent cell surface receptor mediates infection. These same phages each infect subsets of the single CPS-expressing strains, suggesting that some “non-permissive” CPS block access to cell surface receptors, while other “permissive” CPS fail to do so. For phages that do not efficiently infect the acapsular strain, one or more CPS may serve as a direct phage receptor(s) or as a required co-receptor.

To further define the roles of specific CPS during phage infection, we investigated a subset of 6 phages (ARB72, ARB78, ARB82, ARB101, ARB105, and ARB25; marked in blue text in Figure 1). All 6 of these phages infect wild-type *B. thetaiotaomicron* that variably expresses its 8 different CPS and 5 of them (all on Branch 3) infect the acapsular strain poorly or not at all (**Figure 1**). We first tested the hypothesis that some CPS are required as receptors or co-receptors by deleting only the subsets of CPS biosynthetic genes encoding permissive capsules based on our prior experiments with single CPS-expressing strains. For ARB72, which most robustly infects the cps1 and cps3 strains, simultaneous elimination of both of these capsules from wild-type *B. thetaiotaomicron* reduced infection below the limit of detection (**Figure 2A**). Likewise, elimination of the most permissive CPS for the four other Branch 3 phages (ARB78, ARB82, ARB101 and ARB105) significantly reduced *B. thetaiotaomicron* infection by these phages, in some cases in the presence of permissive CPS (**Figure 2B-E**).

**Figure 2.**
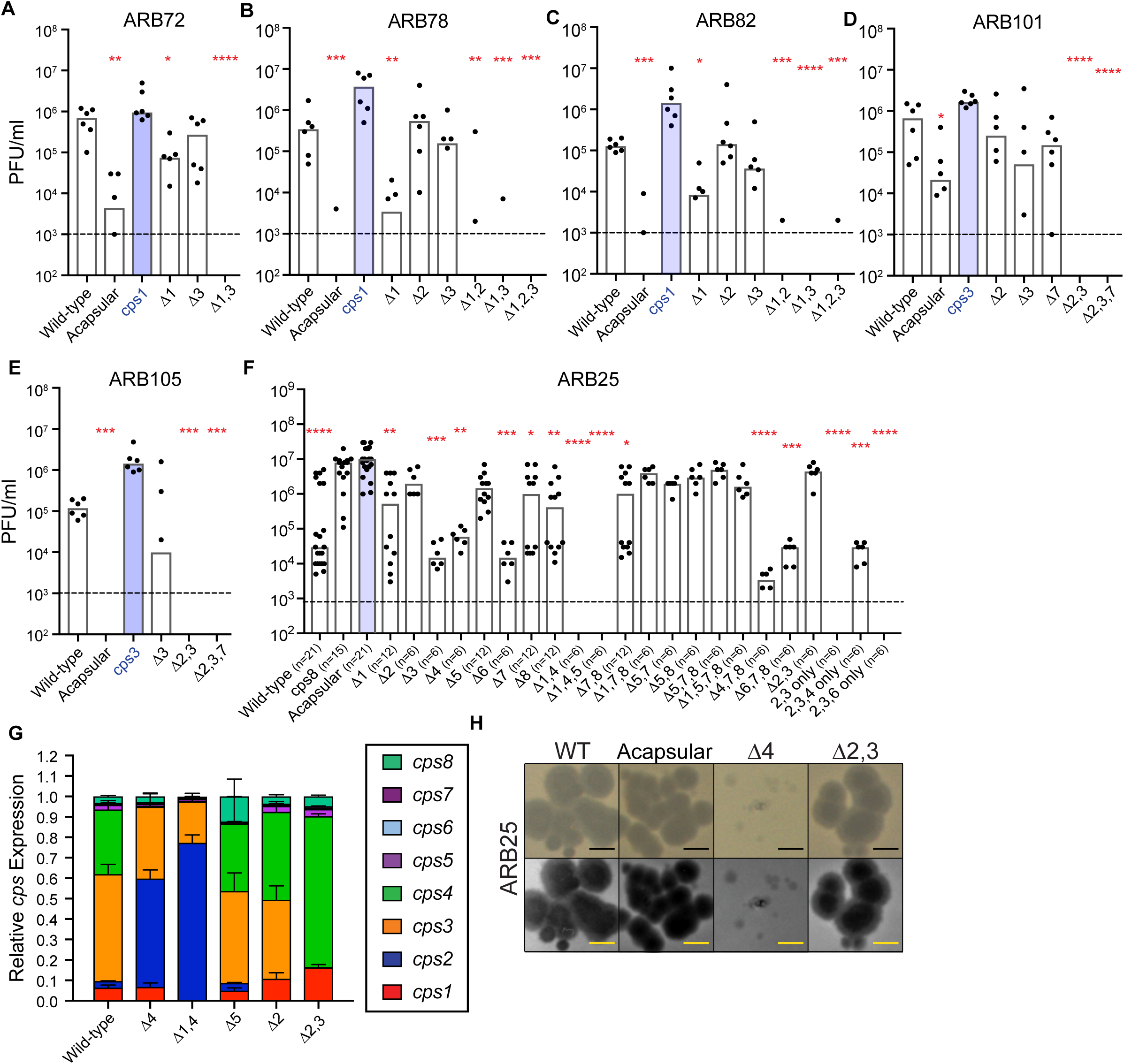
Infection of various CPS mutant strains by Branch 3 phages ARB72 (A), ARB78 (B), ARB82 (C), ARB101 (D) and ARB105 (E) is inhibited by eliminating most or all of the permissive CPS from wild-type *B. thetaiotaomicron*. Each phage was tested on the wild-type strain, the acapsular strain, their respective preferred host strain (blue bars), and a set of bacterial strains harboring selected *cps* locus deletions that correspond to their predetermined host range (n = 6 replicates/phage). (F) Elimination of permissive CPS from Branch 2 phage ARB25 reduces infection, but complete reduction of infection only occurs in the context of deleting more than one permissive CPS. The number of replicates (n=6-21) conducted on each strain is annotated in parentheses next to the strain name. (G) Relative *cps* locus expression of the 8 *cps* loci in the indicated strains. (H) Representative pictures of phage plaques on the indicated host strains. The top row of images for each phage is unaltered; background and unnaturally saturated pixels were removed from images in in the bottom row to facilitate plaque visualization. Scale bar = 2mm. For panels A-F, significant differences in phage titers on the preferred host strain were calculated via Kruskal Wallis test followed by Dunn’s multiple comparisons test. * p < 0.05; ** p < 0.01; *** p < 0.001, **** p < 0.0001. For panel G, significant changes in *cps2* expression were observed in Δ4 and Δ1,4 strains (p < 0.05 for each change, determined by Dirichlet regression). In panels A-F, bars are drawn at the median and individual points shown. In panel G, bars represent mean and error bars SEM, n=3.

For ARB25, which infects 7 of the 10 strains tested in our initial plaque assays (**Figure 1**), some single and compounded *cps* deletions significantly reduced infection rates or reduced them below the limit of detection (**Figure 2F**). While individual deletion of four permissive CPS (CPS1,6,7,8) led to partially reduced infection, so did single eliminations of either of two CPS initially determined to be non-permissive (CPS3 and CPS4). Moreover, deletion of non-permissive CPS4 in combination with deleting the *permissive* CPS1 completely eliminated detectable infection suggesting more complicated regulatory interactions, which are known to occur with *Bacteroides* CPS^19, 20^. Interestingly, strains lacking CPS4 or CPS1/CPS4 compensated by significantly increasing relative expression of the non-permissive *cps2* locus, which could contribute to ARB25 resistance (**Figure 2G**).

A strain expressing only two of the non-permissive CPS (CPS2 and CPS3) could not be detectably infected by ARB25 (**Figure 2F**, “2,3 only”). However, a strain expressing CPS2,3,4 regained some susceptibility (**Figure 2F**, “2,3,4 only”), indicating that when CPS4 is present it is capable of mediating some infection by this phage, which is different than the observation made in Figure 1 and is discussed further below. In contrast to sole expression of CPS2 and CPS3 promoting resistance to ARB25, deletion of the *cps2* and *cps3* loci led to dominant expression of *cps1* and *cps4* genes, which increased infection efficiency and led to the production of clearer plaques (**Figure 2F-H**). Additional support for the idea that loss of CPS4 expression alone modifies ARB25 susceptibility comes from plaque morphologies arising from infection of the *Δcps4* strain, which produced smaller and more turbid plaques, demonstrating that when infection does occur it is less productive (**Figure 2H**). Additional experiments with another *B. thetaiotaomicron* strain that encodes homologs of *cps2, cps5* and cps6 support the conclusion that elimination of these permissive capsules reduces, in some cases, bacteria-phage interaction (**Figure S4**). However, the exogenous presence of a stoichiometric excess of non-permissive CPS2 (∼10^9^-fold more than phage) was not able to block the ability of ARB25 to infect the acapsular strain, suggesting that non-permissive CPS do not inhibit phage infection *in trans* (**Figure S5A**).

### *B. thetaiotaomicron* acquires transient resistance to phage infection

Interestingly, we observed that liquid cultures of the various *B. thetaiotaomicron* strains infected with ARB25 or SJC01 did not show evidence of complete lysis after 36 hours of growth, as determined by optical density at 600 nm (OD_600_) (**Figures 3, S6A**). Previous reports demonstrated that *B. fragilis*^28^ and *B. intestinalis*^29^ exhibited transient resistance to phage infection that could be “reset” through removal of the phage from the culture, although the underlying mechanism of this transient resistance was not determined. Based on these observations, we sought to determine if similar transient resistance occurs with *B. thetaiotaomicron* and if this resistance is dependent on CPS.

**Figure 3.**
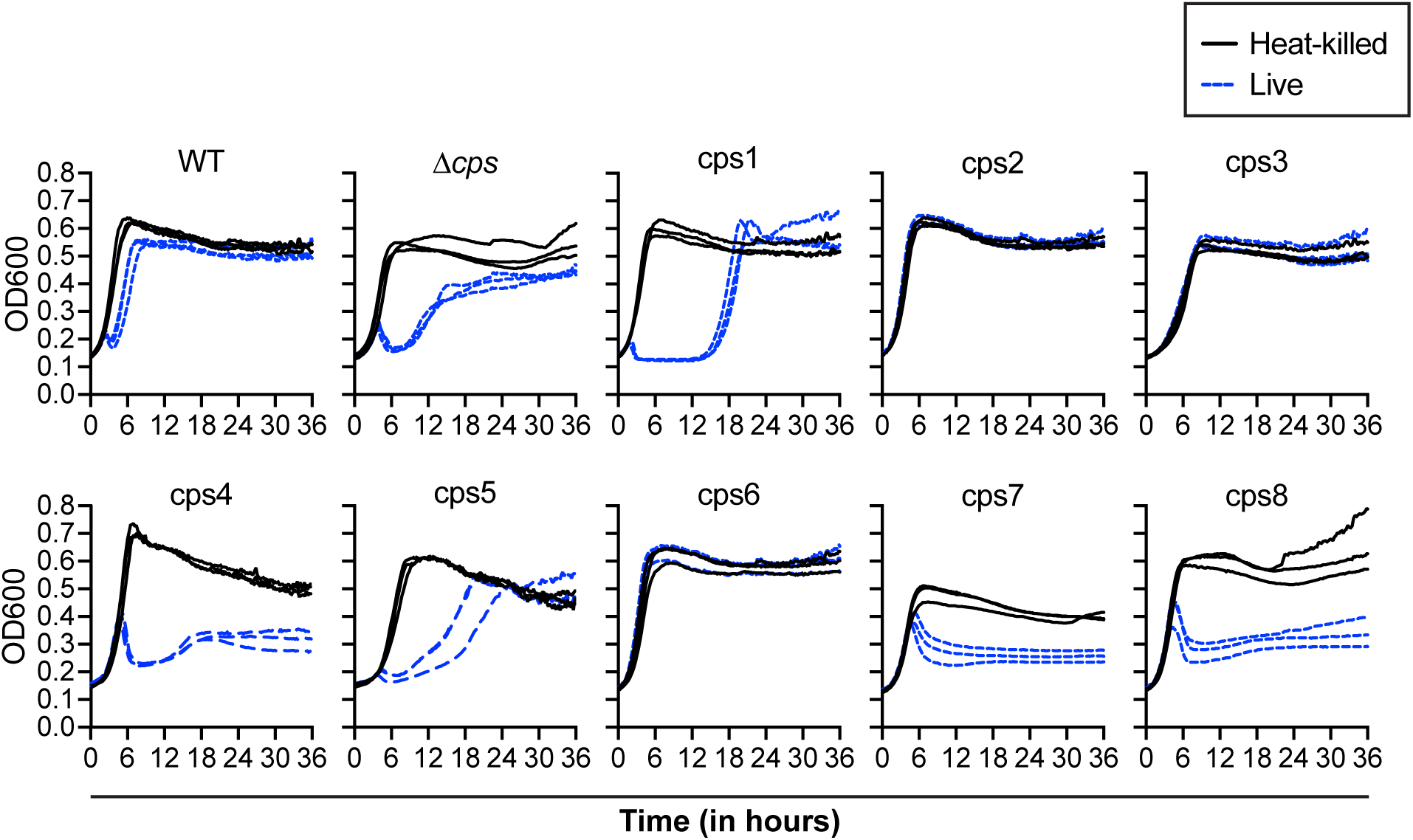
Effects of ARB25 phage infection on growth of bacteria expressing different CPS. Ten strains: the wild-type (WT), the acapsular strain (Δ*cps*), or the eight single CPS-expressing strains were infected with either live or heat-killed ARB25. Growth was monitored via optical density at 600 nm (OD_600_) on an automated plate reading instrument as described in *Methods* and individual growth curves for live and heat-killed phage exposure are shown separately.

Growth curves of each of the CPS-expressing strains inoculated with live or heat-killed ARB25 confirmed our initial host range assays, except, in contrast to the plate-based assays, cultures containing the CPS4-expressing strain were sensitive to killing by this phage in liquid culture, while cultures of the CPS6-expressing strain had no decrease in OD_600_ (**Figure 3**). In these experiments, most strains that appeared to be susceptible via plaque assay exhibited an initial lag in growth or a drop in OD_600_ after growth began. As expected, the *cps*2 and *cps*3 strains did not exhibit any apparent growth perturbation by ARB25. While susceptible strains initiated growth similarly to uninfected cultures and later showed loss of culture density, they subsequently displayed either resumption of growth (wild-type, acapsular, cps1) that approached the density achieved by uninfected controls or growth stagnation at an intermediate culture density (cps4, cps5, cps7, cps8). The former observation suggested outgrowth of a resistant subpopulation of bacteria and culture supernatants taken from ARB25 post-infected, wild-type *B. thetaiotaomicron* still contained high phage titers when exposed to naïve bacteria, excluding the possibility that the phages were inactivated (**Figure S5B**). The observation of growth stagnation after initial loss of bacterial density suggests that a more complex equilibrium is achieved between phage and resistant bacteria that prohibits either from becoming dominant. This behavior was reproducible with 20 separate cultures of the cps4 strain (**Figure S6B**). A similar correlation between plate-based assays and behavior in liquid culture was observed with SJC01, a Branch 2 phage with an infection profile similar to ARB25 (**Figure S6A**). As expected from its resistance to SJC01 in Figure 1, the cps3 strain showed no signs of disrupted growth, whereas cps4, which was non-permissive for both ARB25 and SJC01 in plate-based assays, also showed susceptibility in liquid culture.

We next determined whether strains that had survived or proliferated after exposure to ARB25 retained resistance after removal of phage. In order to isolate phage-free bacterial clones, we isolated individual colonies by sequentially streaking each twice from a subset of the cultures that gained resistance to ARB25 (WT, acapsular, cps1 and cps4) as well as the inherently ARB25-resistant cps2 strain. The majority of clones isolated using this process were free from detectable phage (see *Methods*). We then re-infected each clone with live ARB25 and monitored susceptibility by delayed growth or drop in the culture density as compared to infection with heat-killed phage. As expected, the cps2 strain remained resistant. On the other hand, the majority of clones (42/61 total, ∼69%) derived from the other four strains regained susceptibility (**Table S2**), suggesting that resistance to this phage is not predominantly caused by a permanent genetic alteration.

### Phage-resistant, wild-type *B. thetaiotaomicron* populations exhibit altered *cps* locus expression

Given that CPS type is correlated with resistance to phage infection, we hypothesized that wild-type *B. thetaiotaomicron* cells that are pre-adapted by expressing non-permissive capsules would be positively selected in the presence of phage. To test this, we infected wild-type *B. thetaiotaomicron* with ARB25 and monitored bacterial growth. Cultures treated with a high multiplicity of infection (MOI ≈1) displayed similar growth kinetics as observed previously, with an apparently resistant population emerging after 3-4 hours (**Figure 4A**). Interestingly, bacterial cultures originating from different single colonies displayed variable growth kinetics, with the growth of one clone barely delayed by treatment with live ARB25. Next, we measured if infection with ARB25 resulted in altered CPS expression by the phage-resistant *B. thetaiotaomicron* population. In support of our hypothesis, *B. thetaiotaomicron* exposed to heat-killed phage predominantly expressed CPS3 and CPS4, which we typically observe *in vitro*. Treatment with live ARB25 resulted in a dramatic loss of *cps*1 and *cps*4 expression with a concomitant increase of expression of the non-permissive *cps*3 locus (**Figure 4B**). While reduction in *cps4* expression did not correlate with increased *cps2* as observed with a *Δcps4* strain (**Figure 2G**), the high abundance of *cps3* expressing bacteria in the initial culture may have enabled ARB25 to select for this population in the hours post-infection. Similar growth and expression phenotypes occurred in cultures treated with a low (≈10^-4^) MOI, but with higher initial bacterial growth before a decline (**Figure S7**). Dirichlet regression (see *Methods*) supported significant expression changes for the *cps1*, *cps3*, and *cps4* loci in response to ARB25 (p < 0.01 for experiments with both low and high MOI). Notably, the most resistant of the three bacterial clones (as evidenced by faster outgrowth post-infection) in each of the two experiments (low and high MOI) exhibited similar *cps* locus expression to the other clones after treatment with live phage, but expressed lower levels of permissive *cps*1 and *cps4* and higher levels of non-permissive *cps*3 in heat-killed phage treatment groups (**Figure S8**). This pre-existing variation in CPS expression may contribute to the ability of some clones to resume growth more rapidly after phage challenge because it was already skewed towards non-permissive CPS.

**Figure 4.**
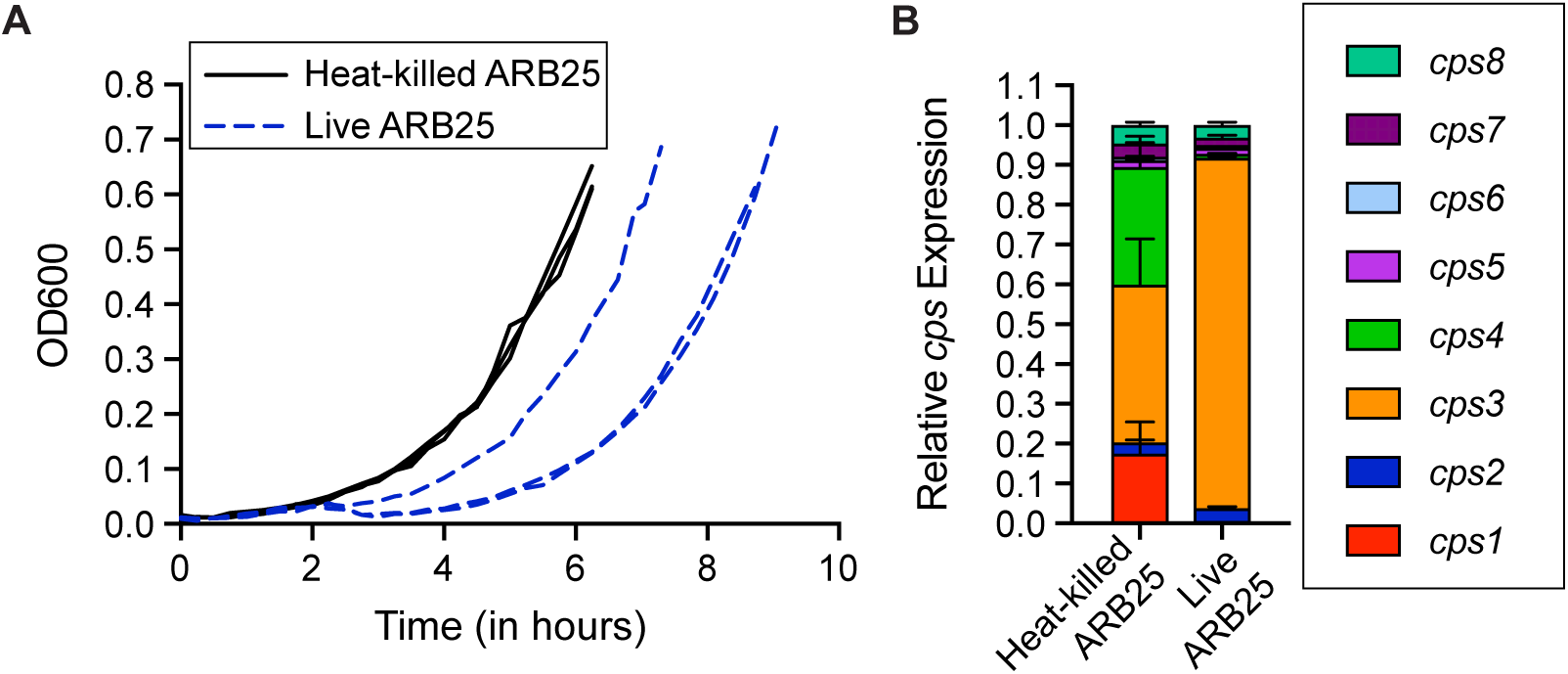
ARB25 infection of wild-type *B. thetaiotaomicron* causes altered *cps* gene expression. Wild-type *B. thetaiotaomicron* was infected with live or heat-killed ARB25 at an MOI of ∼1. (A) Growth was monitored by measuring OD_600_ every 15-30 minutes and individual growth curves for live and heat-killed phage exposure are shown separately (n=3). (B) *cps* gene transcript analysis was carried out by qPCR. The end of the growth curve in panel A represents the point at which cultures were harvested for qPCR analysis (i.e., the first observed time point where culture surpassed OD_600_ of 0.6). Significant changes in *cps1*, *cps3*, and *cps4* expression were observed between groups treated with live or heat-killed ARB25 (p < 0.01 for each, determined by Dirichlet regression; bars represent mean and error bars SEM, n=3). Individual replicates for high and low MOI experiments are displayed in **Fig. S8**.

### Multiple layers of phase-variable featuresequip *B. thetaiotaomicron* to survive phage predation

The results described above support the idea that some individual cells in a *B. thetaiotaomicron* population are pre-adapted to resist phage through expression of different CPS. However, ARB25-infected acapsular *B. thetaiotaomicron* still grew significantly after initial reduction by the phage (**Figure 3**) and most *Δcps*-derived isolates after phage infection had regained susceptibility (**Table S2**), suggesting it is also transient. To determine if additional phage resistance mechanisms are involved, we performed whole genome transcriptional profiling by RNA-sequencing (RNA-seq) to measure transcriptional differences between ARB25 post-infected wild-type and acapsular *B. thetaiotaomicron*. As expected in wild-type, the transcriptional profiles of bacteria surviving after ARB25 infection (n=3) were largely characterized by alterations in CPS expression (**Figure 5A, Table S3a**). Among 83 genes that exhibited significant expression changes ≥3-fold between *B. thetaiotaomicron* exposed to live and heat-killed ARB25, 63 belonged to 4 *cps* loci, with permissive *cps1* and *cps4* decreased and non-permissive *cps2* and *cps3* increased. Interestingly, two additional gene clusters encoding different outer-membrane “Sus-like systems”, which are well-described mechanisms in the Bacteroidetes for import and degradation of carbohydrates and other nutrients^32, 33^, were also decreased in post-infected bacteria. The central features of these systems are outer membrane TonB-dependent transporters (similar to *E. coli* TonA, or **<u>T on</u>**e phage receptor **A**; the first described phage receptor^34^), suggesting the possibility that the proteins encoded by these genes are part of the receptor for ARB25.

**Figure 5.**
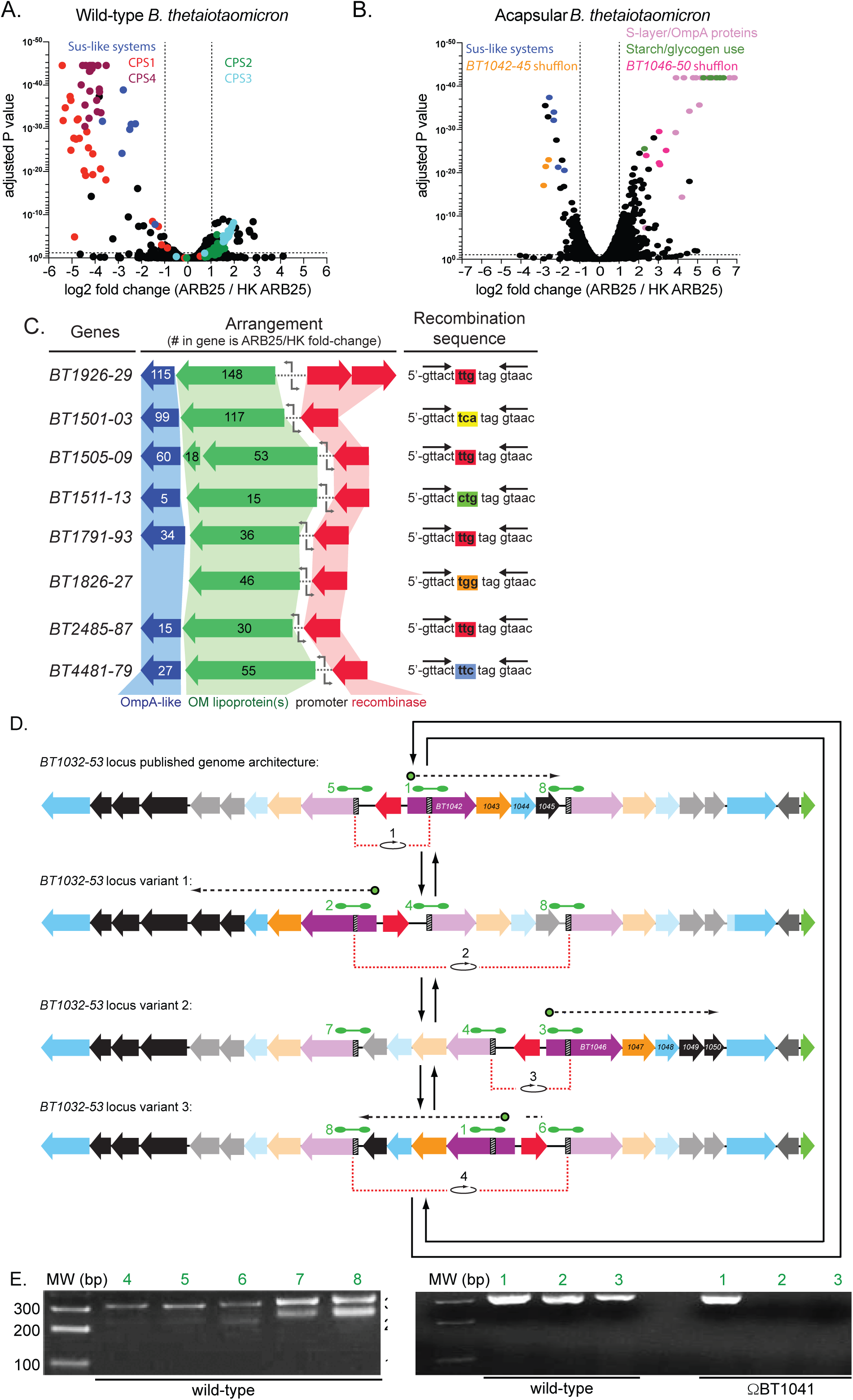
Infection of acapsular *B. thetaiotaomicron* selects for increased expression of multiple phase-variable loci, whereas wild-type mostly alters CPS expression. (A) Wild-type *B. thetaiotaomicron* was infected with ARB25 or was alternatively exposed to heat-killed (HK) ARB25 and cultures were grown to OD_600_=0.6-0.7. Cells were harvested and RNA-seq analysis was carried out as described in *Methods* (n=3 independent experiments for each treatment group). Transcript abundance was compared between live and HK treatments to generate fold change (x axis), which is plotted against the adjusted P value (EdgeR) for each gene. (B) Acapsular *B. thetaiotaomicron* was treated with ARB25 or HK ARB25 and fold change in transcript abundance was calculated, as described in panel A. (C) Among the genes with increased expression in post-infected acapsular *B. thetaiotaomicron*, 25 genes were part of 8 different gene clusters that encode predicted tyrosine recombinases along with outer membrane lipoproteins and OmpA-like proteins. These gene clusters are shown. The number inside the schematic for each gene represents the fold change in expression in ARB25-treated cells relative to those treated with HK ARB25. Flanking the promoters of each of these loci are pairs of imperfect, 17 nucleotide palindromic repeats. PCR analysis and amplicon sequencing of each orientation of these 8 promoters revealed expected confirmation of changes in orientation to the “ON” position in ARB25-exposed acapsular *B. thetaiotaomicron*, although we were unable to quantify the on/off ratios due to high levels of sequence similarity between the 8 loci. (D) Another chromosomal locus with signatures of phage-selected recombination was identified by RNA-seq. Specifically, 3 of 4 genes in an operon (*BT1042-BT1045*) were significantly down-regulated after exposure to phage and 5 genes in an adjacent operon (*BT1046-BT1051*) were up-regulated. (E) PCR using oligonucleotides flanking direct repeats within the *BT1032-BT1053* locus (green dumbbells, panel D) were used to demonstrate locus architecture in wild type *B. thetaiotaomicron* and in a mutant lacking the tyrosine recombinase within this locus (*B. thetaiotaomicron* ΔBT1041). All RNAseq data is provided in **Table S3a-g**.

In the transcriptome of acapsular *B. thetaiotaomicron* subjected to the same live and heat-killed ARB25 treatments described above, 118 genes showed significant expression changes and most of these (100, 85%) were upregulated (**Figure 5B, Table S3b**). One of the two Sus-like systems (*BT2170-73*) that was decreased in ARB25-exposed wild-type was also decreased to similar levels in acapsular *B. thetaiotaomicron*. Among the most highly upregulated genes (28 genes with ≥10-fold increase and an adjusted p-value ≤ 0.01) after ARB25 infection, 6 genes in the well-characterized starch-utilization system (Sus)^32^ were increased in post-infected cultures, suggesting that surviving bacteria are exposed to and metabolize glycogen that is released from lysed siblings. An additional 17 genes belong to 8 loci that encode predicted outer membrane S-layer lipoproteins and OmpA *β*-barrel proteins. One of these (BT1927) was previously investigated and found to be phase-variable and increase *B. thetaiotaomicron* resistance to complement-mediated killing when locked in the “on” state^35^. The remaining S-layer clusters share both syntenic organization and homology to this original S-layer gene cluster. Closer scrutiny of the promoter regions upstream of the newly identified loci revealed that each is also flanked by a pair of imperfect, 17 nucleotide palindromic repeats (**Figure 5C**). Three of these repeats are identical to the repeats known to mediate recombination at the *BT1927* promoter^35^. The remaining 4 sequences only varied by the sequence of a trinucleotide located in the middle of each imperfect palindrome (**Figure 5C**). Finally, amplicon sequencing of the promoter regions using directionally-oriented primers supported the existence of the proposed recombination events in 5 of the 7 newly identified loci, while two did not generate PCR products (**Figure S9**).

Among the remaining genes that were significantly up- or down-regulated in post ARB25-infected acapsular *B. thetaiotaomicron*, there was an additional signature of genes for which DNA recombination is involved in re-organizing expression of cell surface proteins. Specifically, the expression of 3 of 4 genes in an operon (*BT1042-45*) involved in utilization of host *N*-linked glycans^36, 37^ were expressed an average of 4.9-fold less in ARB25-infected acapsular cells. Correspondingly, 5 genes in an adjacent operon (*BT1046-50*) with similar arrangement and predicted functions exhibited an average of 11.9-fold increased expression. Both of these operons have been previously linked to transcriptional regulation by a nearby extra-cytoplasmic function sigma (ECF-*σ*), anti-*σ* factor pair, such that when the single ECF-*σ*coding gene (*BT1053*) is deleted, the ability to activate the adjacent operons is eliminated^16^. Based on 1) the ARB25-dependent shift in gene expression described above; 2) the observation that two genes encoding TonB-dependent transporters (*BT1040, BT1046*) appear to be truncated at their 5’ ends compared to *BT1042* (**Figure 5D**, **S10A**) and only the full-length BT1042 sequence harbors a required anti-*σ* contact domain^16^; and 3) the presence of a gene encoding a putative tyrosine recombinase (*BT1041*) located in the middle of this locus, we hypothesized that this gene cluster possesses the ability to undergo recombination at sites within the three TonB-dependent transporter genes and that specific combinatorial variants are selected under phage pressure.

To test this, we designed PCR primer pairs (**Figure 5D**, green dumbbells) to detect both the originally annotated sequence orientation and 3 potential alternative recombination states derived from either moving the full-length 5’ end of *BT1042* to one of two alternative *susC*-like genes or an internal rearrangement derived from recombination of two incomplete *susC*-like genes (**Figure 5D**, variants 1-3). In support of our hypothesis, we were able to detect by both PCR (**Figure 5E**) and amplicon sequencing (**Figure S10B**) the presence of all 5 predicted alternative recombination states (**Figure 5D,E**), plus the 3 expected from the originally published genome assembly^14^. In further support of our hypothesis, an insertion mutation in the associated tyrosine recombinase-coding gene (*BT1041*) locked the corresponding mutant into the native genomic architecture (**Figure 5E**). Further sequence analysis and tracking of single nucleotide polymorphisms in the 5’ ends of the three recombinationally active *susC*-like genes narrowed the recombination site down to a 7 bp sequence that is flanked by an imperfect direct repeat (**Figure S10B**). Thus, three separate operons that are under the transcriptional control of a single ECF-*σ* regulator and are involved in utilization of host *N*-linked glycans, also undergo recombinational shuffling. This strategy is similar to recombinational shufflons involving nutrient utilization functions in *B. fragilis* ^18, 38^. One explanation is that these shufflons have evolved to subvert phage infection by expressing alternate cell surface receptors that are involved in importing key nutrients but are also targeted by phages. However, elimination of the genes spanning BT1033-52 did not eliminate ARB25 infection in the acapsular strain, suggesting that an additional or different receptor(s) exists. Interestingly, the BT1033-52 mutant exhibited variable plaquing efficiency compared to the acapsular parent (**Figure S11**), suggesting that loss of these genes might exert global effects that mediate susceptibility to ARB25.

To further understand the transcriptional response of *B. thetaiotaomicron* to phage infection, we performed additional RNAseq experiments with ARB25 and the cps1 strain or with a different Branch 2 phage, SJC01. Interestingly, the cps1 strain that is forced to express a permissive capsule can also survive ARB25 infection (**Figure 3**) and mainly does so with transcriptome alterations that involve increased expression of the S-layer proteins identified above (**Figure S12A, Table S3c**). This suggests that, at least for CPS1, co-expression of capsule and S-layer proteins may not be mutually exclusive. Wild-type bacteria infected with SJC01 exhibited similar alterations in CPS expression as were seen with ARB25 (**Figure S12B, Table S3d**). In wild-type bacteria infected with SJC01, 61 of 67 differentially expressed genes belonged to CPS1 and CPS4 (both downregulated) or CPS3 (upregulated), which is consistent with the latter capsule being non-permissive for SJC01. As observed with ARB25, nutrient-utilizing Sus-like systems were also down-regulated in SJC01-infected cells, including previously described systems for ribose^39^ and fungal cell wall *α*-mannan utilization^40^; notably, these systems are different than those down-regulated in ARB25-exposed cultures. Finally, in SJC01-infected acapsular cultures, expression of 4 of the 8 S-layer proteins was prominent, with the *BT1927-25* locus being the most highly expressed feature (**Figure S12C, Table S3e**). An interesting feature of this transcriptome was up-regulation between 6-16 fold of 2 genes (*BT4014-13*) encoding predicted restriction endonucleases (**Figure S12D**). Closer examination of this locus revealed a recombinase located upstream of *BT4014* and predicted 18 bp inverted repeats flanking a near consensus promoter sequence that is oriented away from *BT4014* (*i.e*., “off”) in the assembled genome. To test if this promoter undergoes DNA inversion, we designed primers flanking the predicted recombination sites and performed PCR followed by amplicon sequencing, which confirmed that expression of these genes is also under phase-variable control (**Figure S12E**).

### S-layer expression promotes resistance to multiple phages and is a prominent evasion strategy *in vivo*

The gene encoding the canonical outer membrane S-layer protein (*BT1927*), and its downstream genes (*BT1926-25*) were among the most highly activated in acapsular *B. thetaiotaomicron* that had survived infection with either ARB25 or SJC01 (**Figures 5B, S12C**). We therefore focused on the effects of these proteins on phage infection. Expression of these genes can be locked into the “on” or “off” orientations by mutating the recombination site upstream of the phase-variable promoter^35^ and we re-engineered acapsular *B. thetaiotaomicron* into these 2 expression states. Consistent with the hypothesis that the BT1927 S-layer promotes phage resistance, acapsular S-layer “off” cells were more effectively inhibited by the presence of live ARB25 relative to acapsular S-layer “on” cells (**Figure 6A**). The strength of this effect was altered by the age of the colonies used for subsequent liquid culture experiments to test phage infectivity (**Figure S13**), suggesting that other environmental factors alter the expression or function of this S-layer. Testing of acapsular *BT1927* on/off strains that had been grown under optimal conditions for ARB25 resistance revealed that constitutive expression of the BT1927 S-layer promotes resistance to 3 additional phages from Branches 1 and 2 (**Figure 6B-D**). This latter observation, in combination with previous findings that this S-layer promotes complement resistance^35^, suggests that BT1927 and perhaps the 7 other *B. thetaiotaomicron* S-layers discovered here more broadly promote resistance to a variety of perturbations.

**Figure 6.**
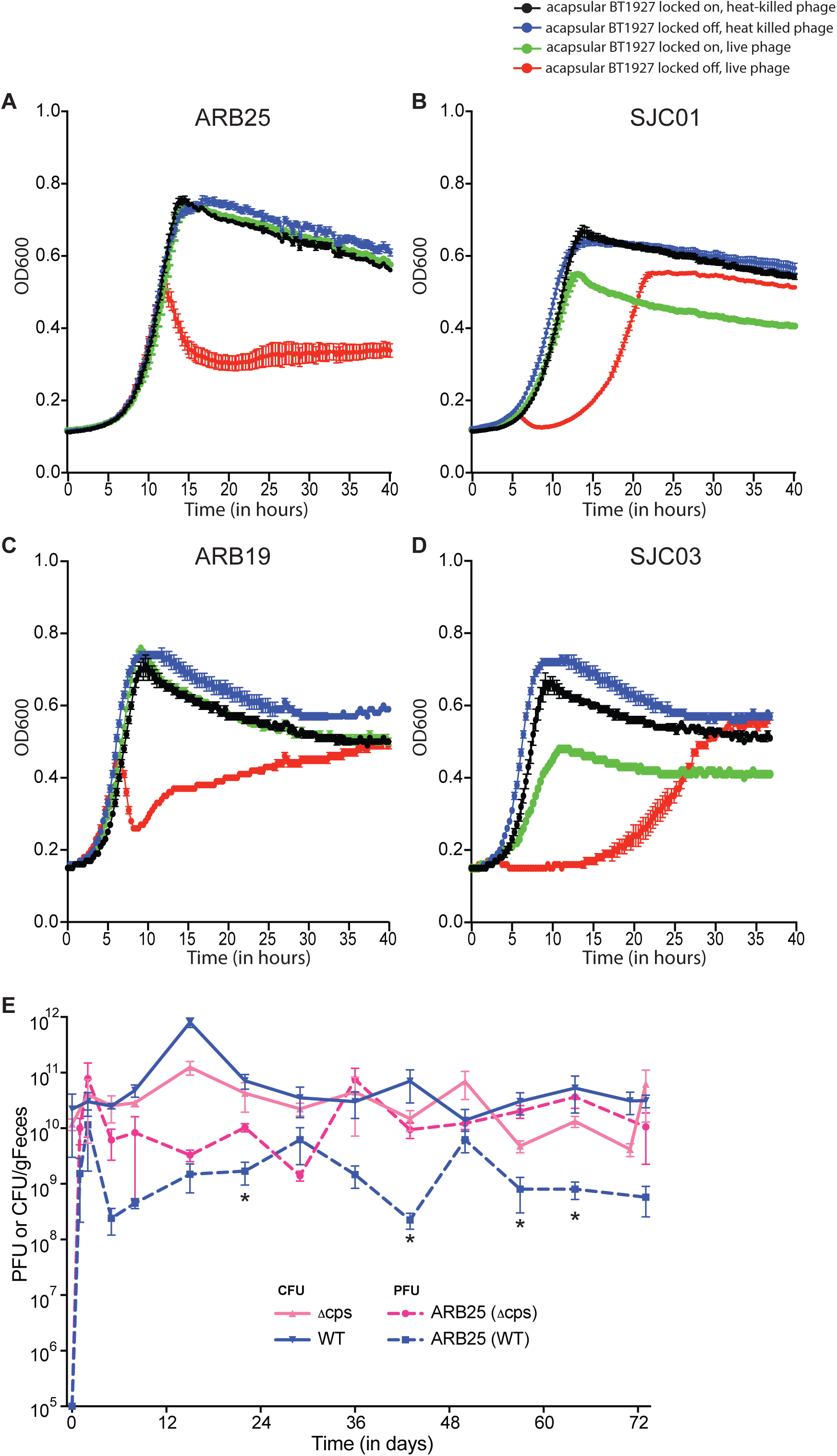
Expression of the BT1927 S-layer increases *B. thetaiotaomicron* resistance to four different phages. Acapsular *B. thetaiotaomicron* S-layer ‘ON’ and ‘OFF’ mutants (Δ*cps* BT1927-ON and Δ*cps* BT1927-OFF, respectively) were infected with (A) ARB25, (B) SJC01 C) ARB19, or D) SJC03 in liquid culture. Growth was monitored via optical density at 600 nm (OD_600_) as described in *Methods* and individual growth curves for live and heat-killed phage exposure are shown separately (n=3-6 per panel). (E) CFU (solid line) or PFU (dashed line) per g Feces for each monocolonized Germ-free swiss webster mouse with either wild-type (blue) or acapsular (Δcps; pink) *B. thetaiotaomicron* and challenged with ARB25 phage (*, p-value ≤ 0.05 student’s t test).

Based on our results, phase-variation of CPS, S-layers, nutrient receptors, and restriction endonucleases are all selected for during phage predation. However, these same phase-variable evasion mechanisms could also explain the previously described observation that *Bacteroides* can co-exist with phage *in vitro* in the phenomenon termed “pseudolysogeny” ^28, 41^. We hypothesized that if the phase-variable systems that promote resistance also spontaneously revert some cells to a susceptible state, then the population could remain mostly resistant but generate a enough susceptible bacteria to continuously propagate phage. To test this in an *in vivo* model with high bacterial density and other features of the colonic environment, we colonized germfree Swiss Webster mice separately with either wild-type or acapsular *B. thetaiotaomicron* for 7 days, then introduced ARB25 by oral gavage. As expected, both bacterial populations reached a high colonization level within 1 day, which was not noticeably perturbed upon addition of phage (**Figure 6E**). ARB25 levels also rose to a high level shortly after introduction and both host bacteria and phage remained present for more than 70 days of co-colonization. Interestingly, while ARB25 levels initially dropped at least 10-fold in all mice between 2 and 4 days after introduction and remained lower than the number of colonizing bacteria, these trends diverged after 30 days with ARB25 levels becoming significantly higher at several timepoints in *Δcps*.

Because it lacks the ability to evade ARB25 through alterations in CPS expression, we further hypothesized that the *Δcps* strain might be more prone to accrue mutations that promote full resistance after several weeks of constant ARB25 pressure. To address this, we isolated *B. thetaiotaomicron* clones from feces and cecal contents of each of the colonized mice and subjected them to serial passage to ensure they were phage-free (see *Methods*). We then measured the susceptibility of these isolates to ARB25 after 10 days of repeated, daily passage *in vitro* (**Table S4**). Contrary to our hypothesis, all of the *Δcps* clones had regained susceptibility to ARB25, while 5/13 wild-type isolates remained resistant, suggesting that longer-term (perhaps permanent) resistance can occur after prolonged exposure to ARB25 *in vivo*, but this requires the presence of CPS.

Lastly, to determine which of the phase-variable resistance mechanisms are operative *in vivo* after prolonged ARB25 exposure, we performed RNAseq on bacteria recovered from the cecal contents of mice after 72 days of phage infection. Compared to the respective wild-type and acapsular strains grown *in vitro* and exposed to heat-killed ARB25, many genes were induced *in vivo* as expected based on previous studies of *B. thetaiotaomicron* adaptation to the diet and host derived nutrient conditions in the gut^36, 42, 43^. Wild-type *B. thetaiotaomicron* that had co-existed with ARB25 for 72d *in vivo* surprisingly exhibited lower expression of non-permissive CPS3 (average 5-fold lower than uninfected wild-type grown *in vitro*). While expression of non-permissive CPS2 was increased, so was expression of several permissive CPS (CPS5, CPS6, CPS7 and CPS8, **Figure S12F**, **Table S3F**), which may have been influenced by growth *in vivo*, which we have previously shown to be selective for *B. thetaiotaomicron* expressing CPS4, CPS5 and CPS6^8^. While wild-type bacteria did not display dominant expression of non-permissive CPS *in vivo* like they did in short-term infection experiments *in vitro*, they did increase expression of 6 of the S-layer loci, while repressing one. Finally, the phase-variable restriction endonuclease system identified in vitro in SJC01 infected cells was also upregulated in wild-type and acapsular *B. thetaiotaomicron* after prolonged co-existence with ARB25 *in vivo* (**Figure S12FG**).

As expected, acapsular *B. thetaiotaomicron* exhibited expression of some of its S-layers after 72 days of ARB25 exposure *in vivo* (**Figure S12G, Table S3g**). However, most of the 8 S-layers only showed modestly increased or reduced expression relative to *in vitro* grown *B. thetaiotaomicron* after prolonged *in vivo* existence with ARB25 and just one of the S-layers (BT1826) became dominant with 2,738-fold increased expression. This latter result suggests that this protein may confer optimal ARB25 resistance in this particular strain background and *in vivo* growth condition. Surprisingly, acapsular *B. thetaiotaomicron* displayed high expression of another set of 3 genes (*BT0292-94;* increased 79-156-fold) and one of these genes BT0294 encodes a predicted lipoprotein of less than ∼50% of the deduced size of the S-layer proteins. Adjacent to this locus is a predicted recombinase. We were able to identify a near-consensus promoter, assembled in the “off” orientation in the *B. thetaiotaomicron* genome, flanked by 18 bp repeats (**Figure S12D**). As with the newly identified S-layer proteins and restriction enzyme system, combined PCR/sequencing demonstrated that this promoter is capable of undergoing phase-variation (**Figure S12E)** bringing the total number of *B. thetaiotaomicron* phase-variable features that show selection in response to phages to 19 members of 4 different functional groups (CPS, S-layers, TonB-dependent transporters and restriction enzymes). Based on the results described above, it is probable that this complex set of phase-variable functions equips *B. thetaiotaomicron* with the versatility to optimally resist phage pressure, but also simultaneously adapt to other environmental conditions such as nutrients and host immunity.

## Discussion

Production of multiple phase-variable CPS is a hallmark of human gut Bacteroidetes (**Figure S1A**). Previous work has revealed the importance of *Bacteroides* CPS in interactions with the host immune system^7, 8, 44, 45^. However, other biological roles for *Bacteroides* CPS have remained unexplored. Using a panel of *B. thetaiotaomicron* strains that express individual CPS, we tested a previously inaccessible hypothesis: that *Bacteroides*-targeting phage can be both inhibited and assisted by the repertoire of CPS expressed by their host bacteria. Our data clearly indicate that production of specific CPS is associated with alterations in phage susceptibility, which is underscored by the observation that none of the 71 phages characterized here infect every CPS-variant that we tested (**Figure 1**). Phage-mediated selection and interactions with the host immune system help to explain both the extensive diversification of CPS structures in gut-resident Bacteroidetes^8, 11^ and their complex phase-variable regulation within a given strain or species^20^. Surprisingly, our results also reveal that additional phase-variable functions are expressed by *B. thetaiotaomicron* during selection by phage, highlighting the diversity of strategies in *Bacteroides* for surviving phage predation.

There are several mechanisms through which CPS could promote or prevent phage infection^46^. First, CPS may sterically mask surface receptors to block phage binding, although additional specificity determinants must be involved because no individual phage that infects the acapsular strain is blocked by all *B. thetaiotaomicron* CPS. These specificity determinants could be driven by CPS structure (physical depth on the cell surface, polysaccharide charge, permeability) or be actively circumvented by the presence of polysaccharide depolymerases on the phage particles, as has been described in other phage-bacterium systems (e.g., *E. coli* K1 and phiK1-5^47^). Alternatively, certain permissive CPS could serve as obligate receptors^48^ or more generally increase the affinity of a phage for the bacterial cell surface. This latter type of adherence to CPS might increase the likelihood that a phage would contact its receptor by sustained interaction with the extracellular matrix. Some combination of these possibilities is likely to explain the host range infection profile for the majority of the phages in this study. Collectively, our observations provide the foundation for future mechanistic work, which will begin with phage genome sequencing, aimed at understanding the physical and chemical interactions that mediate infection of *B. thetaiotaomicron* and other *Bacteroides* by their phages.

Using ARB25 and SJC01 as representatives from our larger collection, we demonstrate that infection with these single phages does not fully eradicate their target *B. thetaiotaomicron* populations *in vitro* and *in vivo* (**Figures 3, 4, 6, S5-S7, S13**). Similar observations were previously made with ΦCrAss001, a phage that infects *B. intestinalis*^29^. Specifically, though ΦCrAss001 formed plaques on lawns of *B. intestinalis*, it failed to eradicate this bacterium in long-term liquid culture. Given the roles of CPS in mediating *B. thetaiotaomicron*-phage interactions, the outgrowth of a phage-resistant sub-population was especially surprising in the context of acapsular *B. thetaiotaomicron*, revealing the existence of multiple phase-variable surface proteins, at least one of which (*BT1927-26*) confers increased resistance to several phages when constitutively expressed (**Figures 6A-D, S13**).

A previous study measured that only 1:1000 *B. thetaiotaomicron* cells in a phage-free environment express the S-layer encoded by *BT1927*^35^. Given that ARB25 non-permissive CPS can comprise up to 40% of the expressed capsule population (e.g., CPS3 in **Figure 4B**), the rapid emergence of cells expressing alternative CPS could be explained by the pre-existing abundance of non-permissive CPS. Resistant cells expressing S-layer may be less frequent and therefore only emerge after longer periods of phage exposure, such as those that we observed *in vivo*. The original *B. thetaiotaomicron* S-layer study also demonstrated that locking the invertible promoter for the *BT1927* S-layer into the “on” orientation facilitated survival against complement-mediated killing^35^, suggesting the existence of orthogonal roles for this and related proteins. Combined with our data on CPS-mediated phage tropism, our observations that the *BT1927*-encoded S-layer confers resistance to some phages, that 7 other homologous systems are also upregulated after exposure to ARB25, that a shufflon exists that harbors three recombinationally-variable nutrient acquisition operons, and that additional phase-variable restriction enzyme and surface protein systems exist in *B. thetaiotaomicron* collectively reveal that that there are at least 19 independent cellular functions in *B. thetaiotaomicron* that could contribute to surviving phage predation by this species.

Phages are the most abundant biological entities in the gut microbiome^49^ and interest in the roles and identities of these gut-resident viruses is increasing as metagenomic sequencing approaches are unveiling a more comprehensive view of their dynamics during health and disease^21, 22, 25^. Although sequence-based approaches are powerful for describing the phages that are present, they do not generate information on the definitive hosts or the mechanisms of individual bacteria-phage interactions, which are likely to be elaborate. These limitations will prohibit full dissection of the ecological interactions that phage exert on bacterial populations in the gut. The approach taken here of isolating phages for a particular host of interest, with added layers of detail like systematic variation of surface CPS when possible, will be an essential complement to high throughput sequencing studies and will help build a foundation of mechanistic gut bacterium-phage interactions.

This work, which primarily focuses on a single strain of *B. thetaiotaomicron*, points to the existence of a very complex relationship between bacteria and phage in the gut microbiome. Considering the possibilities that these interactions could vary over time, differ by host species/strain, and evolve differently within individuals or regionally distinct global populations, the landscape becomes even more complex. Given the diverse adaptive and counter-adaptive strategies that have apparently evolved in the successful gut symbiont *B. thetaiotaomicron* and its relatives, our findings hold important implications for the use of phages to intentionally alter the composition or function of the gut microbiota. While a cocktail of multiple phages could theoretically be harnessed together to elicit more robust alteration of target populations within a microbiome, the complexity of host tropisms and bacterial countermeasures that exist for *B. thetaiotaomicron* imply that a deliberate selection of complementary phage would be needed. If effective phage cocktails need to be further tailored to individual microbiomes or elicit resistance within individuals or populations similar to antibiotics, the prospects for effective gut microbiome-targeted phage therapy could indeed become very complicated. Given these considerations, our findings contribute an important early step towards building a deep functional understanding of the bacterium-virus interactions that occur in the human gut microbiome.

## Methods

### Bacterial strains and culture conditions

The bacterial strains used in this study are listed in **Table S5**. Frozen stocks of these strains were maintained in 25% glycerol at −80°C and were routinely cultured in an anaerobic chamber or in anaerobic jars (using GasPak EZ anaerobe container system sachets w/indicator, BD) at 37°C in *Bacteroides* Phage Recovery Medium (BPRM), as described previously^50^: per 1 liter of broth, 10 g meat peptone, 10 g casein peptone, 2 g yeast extract, 5 g NaCl, 0.5 g L-cysteine monohydrate, 1.8 g glucose, and 0.12 g MgSO_4_ heptahydrate were added; after autoclaving and cooling to approximately 55 °C, 10 mL of 0.22 µm-filtered hemin solution (0.1% w/v in 0.02% NaOH), 1 mL of 0.22 µm-filtered 0.05 g/mL CaCl_2_ solution, and 25 mL of 0.22µm-filtered 1 M Na_2_CO_3_ solution were added. For BPRM agar plates, 15 g/L agar was added prior to autoclaving and hemin and Na_2_CO_3_ were added as above prior to pouring the plates. For BPRM top agar used in soft agar overlays, 3.5 g/L agar was added prior to autoclaving. Hemin, CaCl_2_, and Na_2_CO_3_ were added to the top agar as above immediately before conducting experiments. Bacterial strains were routinely recovered from the freezer stocks directly onto agar plates of Brain Heart Infusion supplemented with 10% horse blood (Quad Five, Rygate, Montana) (BHI-blood agar; or for the SJC phages used in **Figure 1**, on BPRM agar), grown anaerobically for up to 3 days and a single colony was picked for each bacterial strain, inoculated into 5 mL BPRM, and grown anaerobically overnight to provide the starting culture for experiments (note that for the BT1927 S-layer protein experiment shown in **Figure 6**, 3 days of growth on BPRM medium was determined to promote the greatest phage resistance).

For the experiment described in **Figure 2G**, liquid cultures of *B. thetaiotaomicron* were grown in BPRM using the pyrogallol method as described previously. Briefly, a sterile cotton ball was burned and then pushed midway into the tube, after which 200 µL of saturated NaHCO_3_ and 200 µL of 35% pyrogallol solution were added to the cotton ball. A rubber stopper was used to seal the tubes, and tubes were incubated at 37 °C.

### Bacteriophage isolation from primary wastewater effluent

The bacteriophages described in this study were isolated from primary wastewater effluent from two locations at the Ann Arbor, Michigan Wastewater Treatment Plant and from the San Jose-Santa Clara Regional Wastewater Treatment Facility. After collection, the primary effluent was centrifuged at 5,500 rcf for 10 minutes at room temperature to remove any remaining solids. The supernatant was then sequentially filtered through 0.45 µm and 0.22 µm polyvinylidine fluoride (PVDF) filters to yield “processed primary effluent.” Initial screening for plaques was done using a soft agar overlay method^51^ where processed primary effluent was combined with 1 part overnight culture to 9 parts BPRM top agar and poured onto a BPRM agar plate (e.g. 0.5 mL overnight culture and 4.5 mL BPRM top agar was used for standard circular petri dishes [100 mm x 15 mm]). Soft agar overlays were incubated anaerobically at 37 °C overnight. Phages were successfully isolated using three permutations of this assay (see **Table S1**): (1) Direct plating, where processed primary effluent was directly added to overnight culture prior to plating. (2) Enrichment, where 10 mL processed primary effluent was mixed with 10 mL 2XBPRM and 3 mL exponential phase *B. thetaiotaomicron* culture and grown overnight. The culture was centrifuged at 5500 rcf for 10 minutes and filtered through a 0.22 µm PVDF filter. (3) Size exclusion, where processed primary effluent was concentrated up to 500-fold via 30 or 100 kDa PVDF or polyethersulfone size exclusion columns. Up to 1 mL of processed primary effluent, enrichment, or concentrated processed primary effluent was added to the culture prior to adding BPRM top agar, as described above. To promote a diverse collection of phages, no more than 5 plaques from the same plate were plaque purified and a diversity of plaque morphologies were selected as applicable. When using individual enrichment cultures, only a single plaque was purified.

Single, isolated plaques were picked into 100 µL phage buffer (prepared as an autoclaved solution of 5 ml of 1 M Tris pH 7.5, 5 ml of 1 M MgSO4, 2 g NaCl in 500 ml with ddH_2_O). Phages were successfully plaque purified using one of two methods: (1) a standard full plate method, where the diluted phage samples were combined with *B. thetaiotaomicron* overnight culture and top agar and plated via soft agar overlay as described above or (2) a higher throughput 96-well plate-based method, where serial dilutions were prepared in 96-well plates and 1 µL of each dilution was spotted onto a solidified top agar overlay. This procedure was repeated at least 3 times to purify each phage.

High titer phage stocks were generated by flooding a soft agar overlay plate that yielded a “lacey” pattern of bacterial growth (near confluent lysis). Following overnight incubation of each plate, 5 mL of sterile phage buffer was added to the plate to resuspend the phage. After at least 2 hours of incubation at room temperature, the lysate was spun at 5,500 rcf for 10 minutes to clear debris and then filter sterilized through a 0.22 µm PVDF filter.

### Phylogenetic analysis of human gut Bacteroidetes and enumeration of cps biosynthetic gene clusters

Phylogenetic analysis was performed by creating a core gene alignment using a custom, publicly available software package, cognac, written for R (version 3.6.1) with C++ integration via Rcpp (version 1.0.3)^52^. Briefly, genbank files for the 53 isolates were parsed to extract the amino acid sequences and orthologous genes were identified with cd-hit (version 4.7) requiring at least 70% amino acid identity and ensuring that genes were of similar length^53^. The cd-hit output was parsed and core genes were identified as those present in a single copy in all genomes. Amino acid sequences were concatenated and aligned with MAFFT (v7.310)^53, 54^. The concatenated gene alignment was then used as the input for fastTree (version 2.1.10) to generate an approximate maximum likelihood phylogeny^55^. The tree created from the core genome alignment was then midpoint rooted and visualized using phytools (version 0.6.99) ape (version 5.3) R packages respectively^56, 57^.

To identify *cps* loci within each of the 53 genomes, previously annotated *cps* genes^8^ from the type strains of *B. thetaiotaomicron* VPI-5482 (**Figure S1B**), *B. fragilis* NCTC9343, and *B. vulgatus* ATCC 8482 *cps* loci were used to identify pfam models that correspond to the glycosyl transferases (GTs) they contain, which revealed pfam00534, pfam00535, pfam01755, pfam02397, pfam02485, pfam08759, pfam13439, pfam13477, pfam13579, pfam13692, and pfam14305. In addition, we searched for *upxY* and *upxZ* (pfam02357, pfam13614), and protein tyrosine kinase (PTK; pfam02706.) Genes corresponding to these pfam modules were extracted for the 53 genomes. In addition, because we found that a number of apparent *upxY/Z* homologs, which were in species more divergent than the *Bacteroides* noted above, we performed an additional search for homologous genes using the UpxY/Z amino acid sequences from the 3 species listed above. For this, we searched the Integrated Microbial Genomes (IMG) database IMG genome BLASTp tool and an E-value cutoff of 1e-5. Each *cps* locus was confirmed by visually comparing homologous genes within each gene locus neighborhood. Positive hits for the presence of a *cps* locus were required to contain at least one GT, along with at least one Upx or PTK homolog in the adjacent locus.

### Quantitative host range assays

To accommodate the large number of phage isolates in our collection, we employed a spot titer assay for semi-quantitative comparisons of infectivity on each bacterial strain. High titer phage stocks were prepared on their “preferred host strain,” which was the strain that yielded the highest titer of phages in a pre-screen of phage host range (see **Figure 1, Table S1**). Lysates were then diluted to approximately 10^6^ PFU/mL, were added to the wells of a 96-well plate, and further diluted to 10^5^, 10^4^, and 10^3^ PFU/mL using a multichannel pipettor. One microliter of each of these dilutions was plated onto solidified top agar overlays containing single bacterial strains indicated in each figure. After spots dried, plates were incubated anaerobically for 15-24 hours prior to counting plaques. Phage titers were normalized to the preferred host strain.

### Images of phage plaques

To document the morphologies of plaques formed by the purified phages, two sets of plaque pictures were captured: the first set were taken with a Color QCount Model 530 (Advanced Instruments) with a 0.01 second exposure. Images were cropped to 7.5 mm^2^ but were otherwise unaltered. The second set of images were taken on a ChemiDoc Touch instrument (BioRad) with a 0.5 second exposure. Images were cropped to 7.5 mm^2^ and unnaturally high background pixels were removed (Image Lab, BioRad) to facilitate viewing of the plaques. Both sets of images are shown in **Figure S2**. Plaque images in **Figure S4** were taken on a ChemiDoc Touch instrument (BioRad).

### Incubation of ARB25 phage with extracted CPS

Approximately 50-100 PFU of ARB25 in 50 µL phage buffer were mixed with an equal volume of H_2_O or capsule (2 mg/mL) extracted by the hot water-phenol method (as described in reference ^8^) and incubated at 37 °C for 30 minutes. Samples were then plated on the acapsular strain, and plaques were counted after 15-24 hours anaerobic incubation at 37 °C. Counts from two replicates on the same day were then averaged, and the experiment was performed three times. While the size of individual CPS2 polymers is unknown, an estimate of 1,000 hexose sugars per molecule (180,000 Da) would be 9×10^13^ CPS glycans at 1mg/mL. If the CPS were only 10% pure, incubation with 10^3^ ARB25 PFU/mL was estimated to provide at least 10^9^-fold more CPS glycans than PFU.

### Bacterial growth curves with phages

For growth curve experiments, 3 or more individual colonies of each indicated strain were picked from agar plates and grown overnight in BPRM. Then, for experiments in **Figures 3**, **6**, **S6** and **S13** each clone was diluted 1:100 in fresh BPRM and 100 µL was added to a microtiter plate. 10 µL of approximately 5*10^6^ PFU/mL live or heat-killed phage were added to each well, plates were covered with an optically clear gas-permeable membrane (Diversified Biotech, Boston, MA) and optical density at 600 nm (OD_600_) values were measured using an automated plate reading device (BioTek Instruments). Phages were heat killed by heating to 95 °C for 30 minutes, and heat-killed phage had no detectable PFU/mL with a limit of detection of 100 PFU/mL.

In **Figure S5B**, wild-type *B. thetaiotaomicron* was infected with live or heat-killed ARB25, and bacterial growth was monitored via optical density at 600 nm (OD_600_) on an automated plate reader for 12 hours. At 0, 6.02, 8.36, and 11.7 hours post inoculation, replicate cultures were vortexed in 1:5 volume chloroform, centrifuged at 5,500 rcf at 4 °C for 10 minutes, and the aqueous phase was titered on the acapsular strain. No phages were detected in heat-killed controls.

### Generation of phage-free bacterial isolates and determination of their phage susceptibility

To isolate phage-free bacterial clones from ARB25-infected cultures (**Tables S2, S4**), each culture was plated on a BHI-blood agar plate using the single colony streaking method. Eighteen individual colonies were picked from each plate, and each of these clones was re-isolated on a new BHI-blood agar plate. One colony was picked from each of these secondary plates and was inoculated into 150 μL BPRM broth and incubated anaerobically at 37 °C for 2 days. Only one of the clones (a *cps*4 isolate) failed to grow in liquid media. To determine whether cultures still contained viable phage, 50 µL of each culture was vortexed with 20 µL chloroform, then centrifuged at 5,500 rcf for 10 minutes. 10 µL of the lysate was spotted on BPRM top agar containing naïve acapsular bacteria and was incubated anaerobically overnight at 37 °C. Loss of detectable phage in the twice passaged clones was confirmed for most of the clones (79/89, 89%) by the absence of plaques on the naïve acapsular strain.

To determine whether the resulting phage-free isolates were resistant to ARB25 infection, each culture was diluted 1:100 in fresh BPRM, 100 µL was added to a microtiter plate, and 10 µL of either live or heat-killed ARB25 (approximately 5*10^6^ PFU/mL) was added. Plates were incubated anaerobically at 37 °C for 48 hours, and OD_600_ was measured as described above. Cultures were determined to be susceptible to ARB25 by demonstration of delayed growth or drop in OD_600_ compared to heat-killed controls.

### Measurement of *cps* gene expression

For **Figures 2G**, **4B**, and **S7**, overnight cultures were diluted into fresh BPRM to an OD_600_ of 0.01. For **Figure 4B**, 200 µL of approximately 2 x 10^8^ PFU/mL live phage or heat killed phage were added to 5 mL of the diluted cultures. For **Figure S7**, 200 µL of approximately 2 x 10^5^ PFU/mL live phage or heat killed phage were added to 5 mL of the diluted cultures. Bacterial growth was monitored by measuring OD_600_ every 15-30 minutes using a GENESYS 20 spectrophotometer (Thermo Scientific). Cultures were briefly mixed by hand before each measurement. For determination of relative *cps* gene expression, cultures were grown to OD_600_ 0.6-0.8, centrifuged at 7,700 rcf for 2.5 minutes, the supernatant was decanted, and the pellet was immediately resuspended in 1 mL RNA-Protect (Qiagen). RNA-stabilized cell pellets were stored at −80 °C.

Total RNA was isolated using the RNeasy Mini Kit (Qiagen) then treated with the TURBO DNA-free Kit (Ambion) followed by an additional isolation using the RNeasy Mini Kit. cDNA was synthesized using SuperScript III Reverse Transcriptase (Invitrogen) according to the manufacturer’s instructions using random oligonucleotide primers (Invitrogen). qPCR analyses for *cps* locus expression were performed on a Mastercycler ep realplex instrument (Eppendorf). Expression of each of the 8 *cps* synthesis loci was quantified using primers to a single gene in each locus (primers are listed in **Table S6**) and normalized to a standard curve of DNA from wild-type *B. thetaiotaomicron*. The primers used were selected to target a gene specific to each *cps* locus and were previously validated against the other strains that lack the target *cps* locus for specificity^8^. Relative abundance of each *cps*-sepcific transcript was then calculated for each locus. A custom-made SYBR-based master mix was used for qPCR: 20 µL reactions were made with ThermoPol buffer (New England Biolabs), and contained 2.5 mM MgSO_4_, 0.125 mM dNTPs, 0.25 µM each primer, 0.1 µL of a 100 X stock of SYBR Green I (Lonza), and 500 U Hot Start *Taq* DNA Polymerase (New England Biolabs). 10 ng of cDNA was used for each sample, and samples were run in duplicate. A touchdown protocol with the following cycling conditions was used for all assays: 95 °C for 3 minutes, followed by 40 cycles of 3 seconds at 95 °C, 20 seconds of annealing at a variable temperature, and 20 seconds at 68 °C. The annealing temperature for the first cycle was 58 °C, then dropped one degree each cycle for the subsequent 5 cycles. The annealing temperature for the last 34 cycles was 52 °C. These cycling conditions were followed by a melting curve analysis to determine amplicon purity.

### Transcriptomic analysis of B. thetaiotaomicron after phage infection

Whole genome transcriptional profiling of wild-type and acapsular *B.thetaiotaomicron* infected with live or heat-killed ARB25 or SJC01, or from *in vivo* samples, was conducted using total bacterial RNA that was extracted the same as described above (Qiagen RNAEasy, Turbo DNA-free kit) and then treated with Ribo-Zero rRNA Removal Kit (Illumina Inc.) and concentrated using RNA Clean and Concentrator −5 kit (Zymo Research Corp, Irvine, CA). Sequencing libraries were prepared using TruSeq barcoding adaptors (Illumina Inc.), and 24 samples were multiplexed and sequenced with 50 base pair single end reads in one lane of an Illumina HiSeq instrument at the University of Michigan Sequencing Core. Demultiplexed samples were analyzed using SeqMan NGen and Arraystar software (DNASTAR, Inc.) using EdgeR normalization and >98% sequence identity for read-mapping. Changes in gene expression in response to live ARB25 infection were determined by comparison to the heat-killed reference: retained were genes with ≥ 3-fold expression changes up or down and EdgeR adjusted *P* value ≤ 0.01. All RNA-seq data have been deposited in the publicly available NIH gene expression omnibus (GEO) database as project number GSE147071.

### PCR and sequencing of phase variable B. thetaiotaomicron chromosomal loci

We found that each of the 8 chromosomal loci shown in **Figure 5C** had nearly identical 301 bp promoter sequences, including both of the imperfect palindromes that we predict to mediate recombination and the intervening sequence at each locus. While the 8 S-layer genes and the 7/8 of the upstream regions encoding putative tyrosine recombinases (all but the BT1927 region) shared significant nucleotide identity and gene orientation, we were able to design primers that were specific to regions upstream and downstream of each invertible promoter and used these to generate an amplicon for each locus that spanned the predicted recombination sites. After gel extracting a PCR product of the expected size for each locus, which should contain promoter orientations in both the “on” and “off” orientations, we performed a second PCR using a universal primer that lies within the 301 bp sequence of each phase-variable promoter and extended to unique primers that anneal within each S layer protein encoding gene. Bands of the expected size were excised from agarose gels, purified and sequenced using the primer that anneals within each S layer encoding gene to determine if the predicted recombined “on” promoter orientation is detected. (Note that the assembled *B. thetaiotaomicron* genome architecture places all of these promoters in the proposed “off” orientation. We were able to detect 6/8 of these loci in the “on” orientation in ARB25-treated cells by this method, **Figure S9**). Similar approaches were used to determine the re-orientation of DNA fragments in the *B. thetaiotaomicron* PUL shufflon shown in **Figure 5D** and restriction enzyme and additional lipoprotein system shown in **Figure S12D**. For shufflon gene orientation, we used PCR primer amplicons positioned according to the schematic in **Figure 5D** followed by sequencing with the primer on the “downstream” end of each amplicon according to its position relative to the shuffled promoter sequence. For a list of primers used see **Table S6**.

### Construction of acapsular B. thetaiotaomicron S-layer ‘ON’ and S-layer ‘OFF’ mutants

Acapsular *B. thetaiotaomicron* S-layer ‘ON’ and ‘OFF’ mutants (Δ*cps* BT1927-ON and Δ*cps* BT1927-OFF, respectively) were created using the Δ*tdk* allelic exchange method^58^. To generate homologous regions for allelic exchange, the primers BT_1927_XbaI-DR and BT_1927_SalI-UF were used to amplify the BT1927-ON and BT1927-OFF promoters from the previously-constructed BT1927-ON and BT1927-OFF strains^35^ via colony PCR using Q5 High Fidelity DNA polymerase (New England Biolabs). Candidate Δ*cps* BT1927-ON and Δ*cps* BT1927-OFF mutants were screened and confirmed by PCR using the primer pair BT1927_Diagnostic_R and BT1927_Diagnostic_F and confirmed by Sanger sequencing using these diagnostic primers. All plasmids and primers are listed in **Tables S5** and **S6**, respectively.

### Construction of B. thetaiotaomicron mutants lacking one or more cps loci

All publically available bacterial genomes in NCBI GenBank were queried via MultiGeneBlast^59^ to identify fully sequenced bacteria with *B. thetaiotaomicron* VPI-5482-like cps loci. *B. thetaiotaomicron* 7330 was identified as having VPI-5482-like *cps2*, *cps5*, and *cps6* loci. Mutants of strain 7330 lacking one or more *cps* loci constructed for this study (**Table S5**) were created using the *tdk* allelic exchange method^58^. The *B. thetaiotaomicron* 7330 *tdk*-strain was generated by UV mutagenesis by exposing a liquid culture of 7330 to 320 nm ultraviolet light from a VWR-20E transilluminator (VWR) for 60 seconds and plating onto BHIS-Blood agar supplemented with 200 micrograms/mL of 5-fluoro-2’-deoxyuridine (FUdR). All plasmids and primers used to construct these strains are listed in **Tables S5** and **S6**, respectively.

### Germfree mouse experiments

Germfree Swiss webster mice were gavaged with either wild-type or acapsular *B. thetaiotaomicron* for 7 days of mono-colonization. After 7 days, mice were gavaged with 1M sodium bicarbonate followed immediately by 2×10^8^ PFU of ARB25 as previously described^24^. Feces were monitored for both colony forming units (CFU) or plaque forming units (PFU) every 7 days by plating fecal homogenates in SM buffer or fecal homogenate supernatant and serial dilutions in SM buffer on BPRM top agar plates.

### Data representation and statistical analysis

The heatmaps for **Figures 1** and **S3** and the dendrogram for **Figure 1** were generated in R using the “heatmap” function. Other graphs were created in Prism software (GraphPad Software, Inc., La Jolla, CA). Statistical significance in this work is denoted as follows unless otherwise indicated: * p < 0.05; ** p < 0.01; *** p < 0.001; **** p < 0.0001. Statistical analyses other than Dirichlet regression were performed in Prism. Dirichlet regression was performed in R using the package “DirichletReg” (version 0.6-3), employing the alternative parameterization as used previously^8, 60^. Briefly, the parameters in this distribution are the proportions of relative *cps* gene expression and the total *cps* expression, with *cps7* expression used as a reference since we previously determined this *cps* to be poorly activated and not subject to phase-variable expression^8^. The variable of interest used in **Figure 2G** is bacterial strain, whereas the variable of interest used in **Figure 4B** is phage viability (live versus heat-killed phage). Precision was allowed to vary by group given this model was superior to a model with constant precision, as determined by a likelihood ratio test at significance level p < 0.05.

## Acknowledgements

We thank Rey Honrada at the San Jose-Santa Clara Wastewater Treatment Plant and the staff at the Ann Arbor Wastewater treatment plant for assistance in collecting primary sewage effluent and Dylan Maghini for assistance in identifying shared *cps* loci between *B. thetaiotaomicron* strains VPI-5482 and 7330. This work was funded by NIH grants (GM099513 and DK096023 to E.C.M), an NIH postdoctoral NRSA (5T32AI007328 to A.J.H.), a Stanford University School of Medicine Dean’s Postdoctoral Fellowship (A.J.H.), the NIH Cellular Biotechnology Training Program (N.T.P., T32GM008353) and NIH Bioinformatics Training Grant (R.C., T32GM070449).

## Author Contributions

NTP, AJH, BDM, JOG, JJF, RWPG, and SS performed the experiments. NTP, AJH, JJF and ECM designed the experiments, and analyzed and interpreted the data. RDC and ESS performed whole genome phylogenetic analysis and JJF conducted the corresponding *cps* locus search. JLS and ECM provided tools and reagents. NTP, AJH, JJF, and ECM prepared the display items. NTP, AJH, JJF and ECM wrote the paper. All authors edited and approved the manuscript prior to submission.

---

**Figure S1**. Diversification and structure of *cps* gene clusters in human gut Bacteroidetes. (A) The genomes of 53 different human gut Bacteroidetes (predominantly named type strains) were searched for gene clusters that contain two or more different protein families indicative of *cps* loci (see *Methods*). The number of *cps* loci detected in each genome is shown in the context of phylogenetic tree derived from the core genome of the 53 species used for this analysis; species for which *cps* loci were not detected using our search criteria are marked with a red “X”. Due to gaps in several genomes, which often occur at *cps* loci, the numbers shown are likely to be an underestimate. (B) Schematics of the 8 annotated cps loci in *B. thetaiotaomicron* VPI-5482, which are singly present in the cps1-cps8 strains used in this study, or completely eliminated in the acapsular strain. Genes are color coded according to the key at the bottom and additional Pfam family designations are provided under most genes. The four main protein families used for informatics analysis are marked with asterisks and highlighted in bold in the key.

**Figure S2**. Representative pictures of phage plaques for all phages in this study: (A) phages from Ann Arbor (ARB); (B) phages from San Jose (SJC). The top row of images for each phage are unaltered; background and saturated pixels were removed from images in the bottom row to facilitate viewing of the plaques. Scale bar = 2 mm

**Figure S3**. Replication of a subset of host range assays of *B. thetaiotaomicron*-targeting phages on strains expressing different CPS types. Ten bacteriophages isolated and purified on the wild-type, acapsular, or the 8 single CPS-expressing strains were re-tested in a spot titer assay to determine phage host range. 10-fold serial dilutions of each phage ranging from approximately 10^6^ to 10^3^ plaque-forming units (PFU) / ml were spotted onto top agar plates containing the 10 bacterial strains. Plates were then grown overnight, and phage titers were calculated. Titers are normalized to the titer on the preferred host strain for each replicate. Each row in the heatmap corresponds to a replicate for an individual phage, whereas each column corresponds to one of the 10 host strains. One to three replicates of the assay were conducted for each phage by the two lead authors (AJH and NTP). Assays were carried out at the same time, and each author used the same set of cultures and phage stocks. For comparison, individual replicates from **Figure 1** are included (marked with *).

**Figure S4**. Effects of eliminating permissive CPS from another *B. thetaiotaomicron* strain. (A) We identified *B. thetaiotaomicron* 7330^61^ as the only sequenced and genetically tractable strain that contains VPI-5482-like *cps* loci (*cps2, cps5, and cps6*). We also observed that the Branch 2 phage SJC01did not yield productive infection in *B. thetaiotaomicron* 7330, but could partially clears lawns of *B. thetaiotaomicron* 7330 at high titers. This ability to clear established lawns is a previously described phenomenon known as “lysis from without^62^”. (B) Deletion of permissive capsules (*cps2, cps5*, and *cps6*) either alone or in combination affects VPI-5482 infection by SJC01. (C) Deletion of *B. thetaiotaomicron* VPI-5482-like *cps* loci from *B. thetaiotaomicron* 7330 affects the “lysis from without” phenotype. While SJC01 plaques on WT *B. thetaiotaomicron* VPI-5482, it does not form plaques on wild-type *B. thetaiotaomicron* 7330. However, SJC01 does exhibit a “lysis from without” clearing phenotypes at high densities of phage (top two spots, made with 1 microliter of 1e8 and 1e7 PFU per mL, according to titers observed on wild-type VPI-5482). (D) *B. thetaiotaomicron* 7330 strains lacking *cps5* (with the exception of 7330 Δ*cps5* Δ*cps6*) show the lysis from without phenomenon less frequently than strains that have intact *cps5* (at least n=3 replicates per phage/host pair). For panel B, significant differences in phage titers on each mutant strain were compared to wild type via Mann-Whitney test, * p < 0.05; ** p < 0.01; *** p < 0.001, **** p < 0.0001.

**Figure S5**. Free CPS does not inhibit ARB25 infection when provided *in trans*. (A) ARB25 was incubated with purified CPS1 or CPS2 (1 mg/ml, an estimated 10^9^ molar excess of CPS molecules to phage, see *Methods*) before plating on the acapsular strain, and plaques were counted after overnight incubation. Titers are normalized to mock (H_2_O) treatment. No significant differences in titers were found compared to mock treatment, as determined by Welch’s t test (n=3 biological replicates, bars represent mean ± SEM). (B) Post ARB25-infected, surviving cultures still contain infectious phages. Wild-type *B. thetaiotaomicron* was infected with live or heat-killed ARB25, and bacterial growth was monitored via optical density at 600 nm (OD_600_). At 0, 6.02, 8.36, and 11.7 hours post inoculation, replicate cultures were removed and phage levels were titered (n=3 and individual replicate curves are shown). No phages were detected in heat-killed controls. Note that the PFU/mL do not increase substantially after the initial “burst” corresponding to decreased bacterial culture density prior to re-growth.

**Figure S6**. Effect of CPS and phage infection on bacterial growth. (A) Ten strains: the wild-type (WT), the acapsular strain (Δcps), or the eight single CPS-expressing strains were infected with either live or heat-killed SJC01. (B) 20 different colonies of cps4 or cps5 strains were infected with ARB25. Growth was monitored via optical density at 600 nm (OD_600_) on an automated plate reading instrument as described in *Methods* and individual growth curves for live and heat-killed phage exposure are shown separately.

**Figure S7**. Infection of wild-type *B. thetaiotaomicron* at a low multiplicity of infection and subsequent effects on *cps* gene expression. (A) The wild-type (WT) strain was infected at a low multiplicity of infection (MOI = 1 x 10^-4^) of live or heat-killed ARB25, and bacterial growth was monitored via OD_600_ (n=3 biological replicates and separate curves are shown). (B) RNA was harvested from cultures after reaching an OD_600_ of 0.6-0.7, cDNA was generated, and relative expression of the 8 *cps* loci was determined by qPCR (histogram bars are mean ± SEM of 3 biological replicates. Individual replicates are shown in **Fig. S8**).

**Figure S8**. Single replicates of *cps* expression in heat-killed versus live phage-treated *B. thetaiotaomicron*. Relative *cps* transcript abundance in ARB25 infection experiments at high MOI (A) and low MOI (B). In the high MOI experiment, replicate 2 showed higher starting expression of the non-permissive CPS3 compared to others. In the low MOI experiment, replicate 3 showed higher starting expression of the non-permissive CPS3. In both experiments, post phage-exposed replicates displayed nearly identical CPS expression profiles characterized by high expression of CPS3.

**Figure S9**. Determination of phase-variable promoter switching for six loci encoding putative S-layer proteins. The hypothesis that the promoters associated with seven newly identified *B. thetaiotaomicron* S-layer like lipoproteins was validated using a PCR amplicon sequencing strategy. Because of high nucleotide identity in both the regions flanking the 7 new loci, a nested PCR approach was required to specifically amplify and sequence each site. In the first step, a primer lying in each S-layer gene (**Table S5** “S-layer gene” primers) was oriented towards the promoter and used in a PCR extension to a primer in the upstream recombinase gene (**Table S5** “recombinase gene 3” primer). The products of this PCR were purified without gel extraction and used in a second reaction with a nested primer that lies internal to the previous recombinase gene primer (**Table S5** “recombinase 2” primer). The expected PCR products from this reaction, which are ∼1 kb and span promoter sequences in both the ON and OFF orientations, were excised and used for an orientation-specific PCR using the original S-layer gene primer for each site and a universal primer (green schematic) that was designed for each promoter and is oriented to extend upstream of the S-layer gene (e.g., OFF orientation). Resulting products from this third reaction, which should correspond to the ON orientation if a promoter inversion has occurred in some cells, were obtained for 5/7 of the newly identified loci and the BT1927 S-layer locus as a control. In all cases in which an amplicon and sequence were obtained, the expected recombination occurred between the inverted repeat site proximal to the S-layer gene start (new DNA junction), which would orient the promoter to enable expression of the downstream S-layer gene. The sequences shown are the consensus between forward and reverse reads for each amplicon. The putative core promoter −7 sequence is shown in bold/red text, the coding region of each S-layer gene is shown in bold/blue text and the S-layer gene proximal recombination site is noted and highlighted in bold/gold text. Note that the 5’-end of the sequenced amplicon was not resolved for the BT2486 locus.

**Figure S10**. Recombination between the genes BT1040, BT1042, and BT1046. (A) Pfam domain schematics of the amino acid sequences of these three genes highlighting that BT1040 and BT1046, as originally assembled in the *B. thetaiotaomicron* genome sequence, lack additional N-terminal sequences that are present on BT1042. (B) Sequencing of the 8 PCR amplicons schematized in **Figure 5D**. Amplicons 1, 5 and 8 represent the original genome architecture, while the others represent inferred recombination events that are validated here by sequencing. The 5’ and 3’ ends of the BT1042, BT1040 and BT1046 genes are color-coded to assist in following their connectivity changes after recombination. A series of single-nucleotide polymorphisms (SNPs) present in BT1042, downstream of the proposed recombination site, are highlighted in yellow. The transfer of these SNPs to a fragment containing the 5’ end of BT1040 (Amplicon 4) was used to narrow the recombination region to the 7 nucleotide sequence highlighted in red. Additional SNPs that are specific to the regions upstream of this recombination site are shown in white text for each sequence.

**Figure S11**. The BT1033-52 locus does not affect susceptibility of acapsular *B. thetaiotaomicron* to ARB25. Ten-fold serial dilutions of ARB25 were spotted onto lawns of *B. thetaiotaomicron* Δ*cps* (n=5) and *B. thetaiotaomicron* Δ*cps* ΔBT1033-52 (n=5, n=3 independent clones each with all 15 replicates shown individually). Plaquing efficiency was determined by normalizing plaque counts on *B. thetaiotaomicron* Δ*cps* ΔBT1033-52 relative to plaque counts on *B. thetaiotaomicron* Δ*cps* for each replicate. Statistical significance was determined using the Mann-Whitney test.

**Figure S12**. Whole genome transcriptional analyses of several additional *B. thetaiotaomicron* strain and phage combinations. (A) Infection of the cps1 strain with ARB25, revealing a post-infection response that is largely characterized by increased expression of S-layer/OmpA proteins. (B) Infection of wild-type *B. thetaiotaomicron* with SJC01, revealing that, as with ARB25/wild-type, the bacteria survive phage infection by mostly altering CPS expression. Expression of the non-permissive CPS3 is prominently increased. (C) Infection of acapsular *B. thetaiotaomicron* with SJC01, revealing that in the absence of CPS survival is mostly promoted by increased S-layer/OmpA expression and expression of a newly identified, phase-variable restriction enzyme system. (D) Gene schematic of the newly identified phase-variable restriction enzyme system (top) and a lipoprotein contain locus (bottom) that is different from the 8 S-layer loci also revealed in this study. The inverted repeat sequence that was determined to mediate recombination in each locus is shown. (E) PCR analysis of the restriction enzyme system and new lipoprotein promoter orientations with primers designed to detect phase variation from off to on states. Amplicons were sequenced to confirm the re-orientation to the on orientation (not shown). (F) Global transcriptional responses of wild-type *B. thetaiotaomicron* in the ceca of mice after 72 d of co-existence with ARB25. Note that shifts in CPS expression are mostly characterized by increases in permissive CPS, which may be dictated by growth *in vivo* selecting for these capsules or against the non-permissive CPS3. Correspondingly, wild-type shows increased expression of some but not all S-layer/OmpA systems and the phase-variable restriction enzyme. (G) Global transcriptional responses of acapsular *B. thetaiotaomicron* in the ceca of mice after 72 d of co-existence with ARB25. In the absence of CPS, surviving bacteria show increased expression of only a subset of the identified S-layer/OmpA proteins, with BT1826 expressed most dominantly, along with the newly identified BT0291-94 locus and expression of the restriction enzyme system.

**Figure S13**. ARB25 or SJC01 infection of the acapsular BT1927 locked on and off strains after 1, 2 or 3 days of growth on BPRM. Three separate colonies were picked each day, grown overnight and used to setup infection cultures that were monitored for 24 hours in an automated plate reader. Colonies picked after only 1 day show the least resistance to either phage when BT1927 is locked on. After 2 days, resistance is increased and this continues to increase after 3 days, becoming almost complete (compared to HK controls for ARB25).

**Table S1**. Phages used in this study and details on their isolation.

**Table S2**. Susceptibility of strains to infection by ARB25 after infection and passaging.

**Table S3**. Genes that are differentially regulated in post ARB25-infected wild-type and acapsular *B. thetaiotaomicron*. Consists of sheets a.-g. corresponding to individual RNA-seq comparisons.

**Table S4**. Resistance to ARB25 of wild-type and acapsular *B. thetaiotaomicron* strains after in vivo existence with ARB25 for 72 days and 10 d of repeated culture/passage outside of the mouse.

**Table S5**. Bacterial strains and plasmids used in this study.

**Table S6**. Primers used in this study.

