## Supplementary figures and images for "Multiple phase-variable mechanisms, including capsular polysaccharides, modify bacteriophage susceptibility in *Bacteroides thetaiotaomicron*"

### Supplemental Figure 1

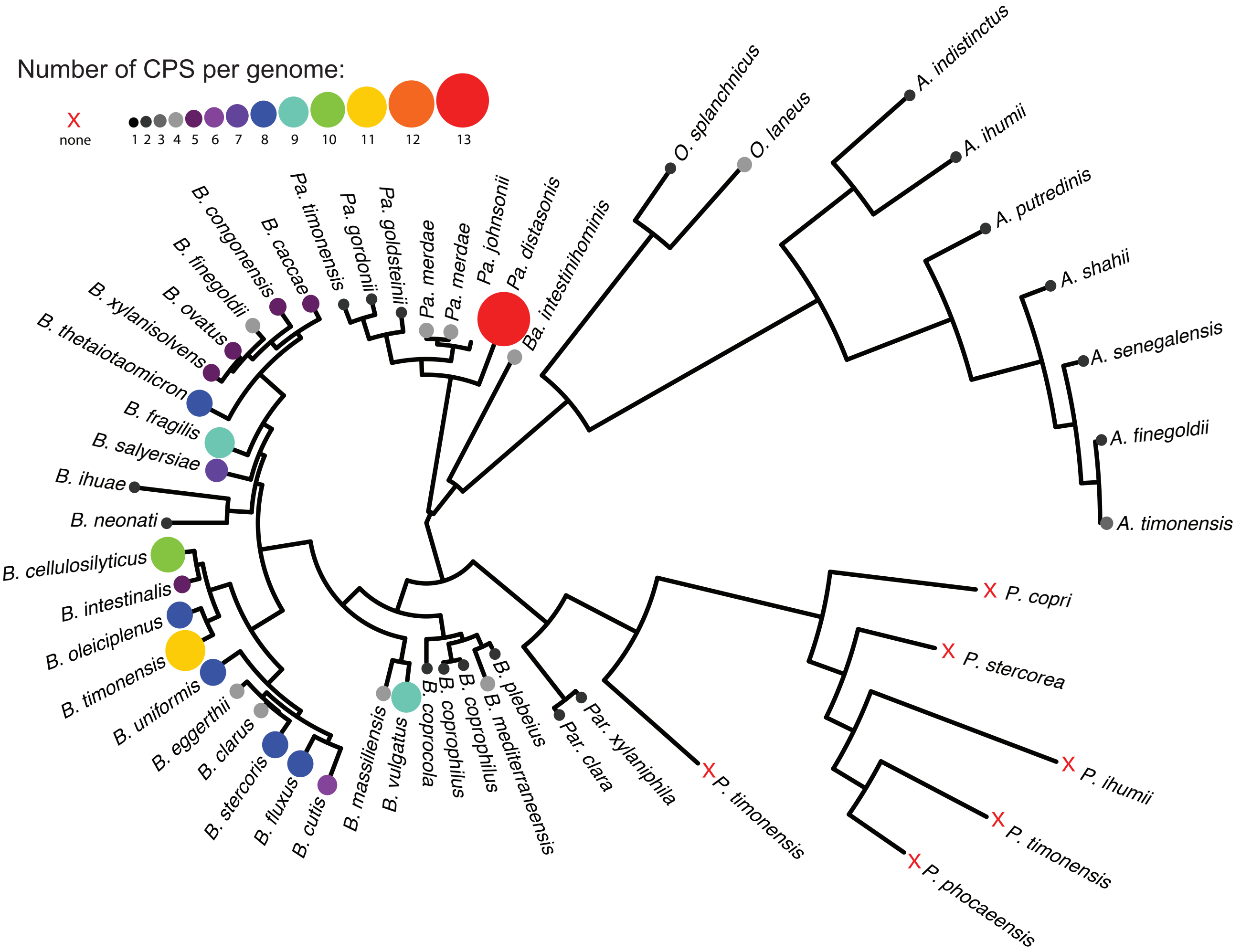

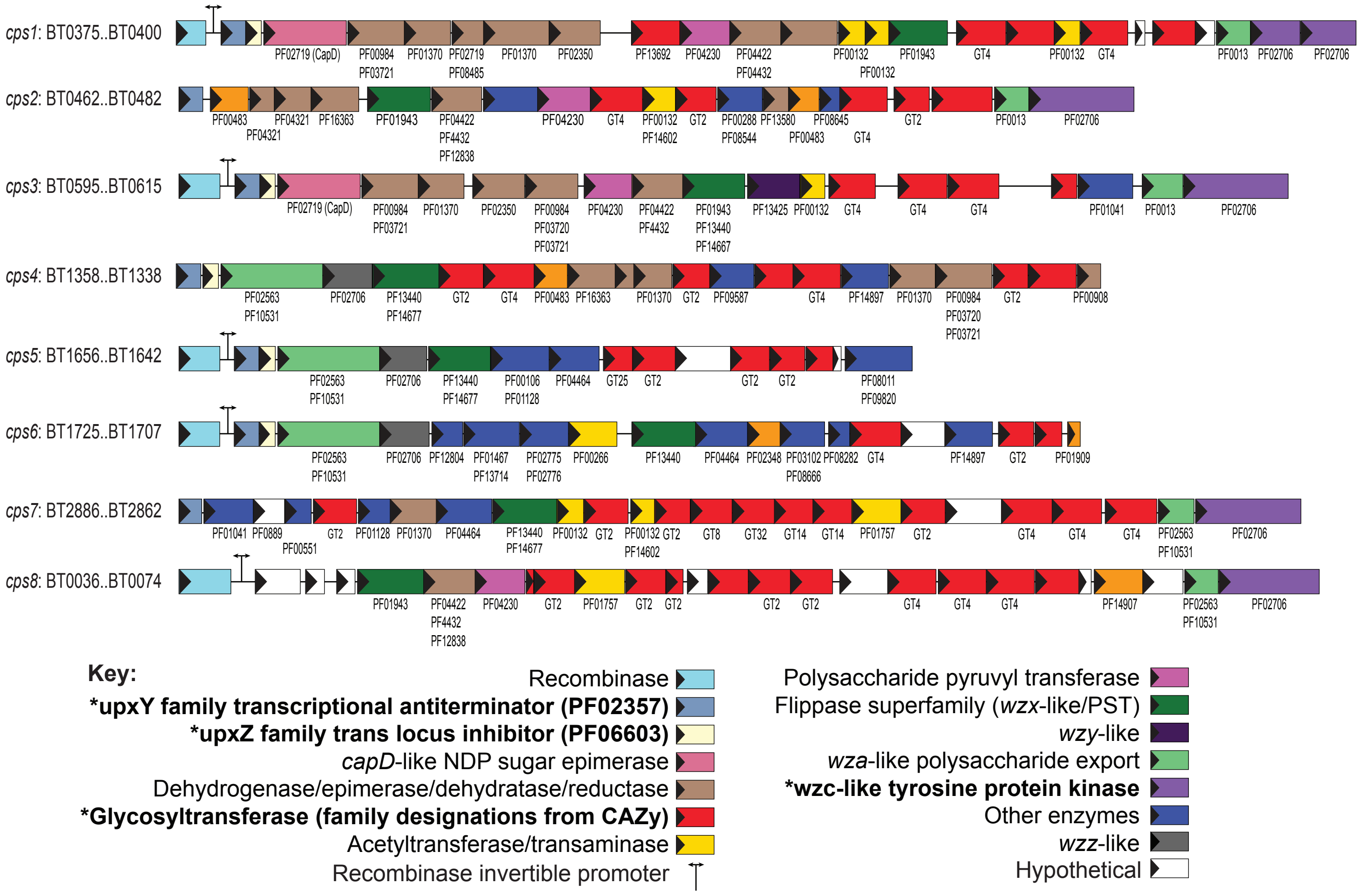

### Supplemental Figure 2

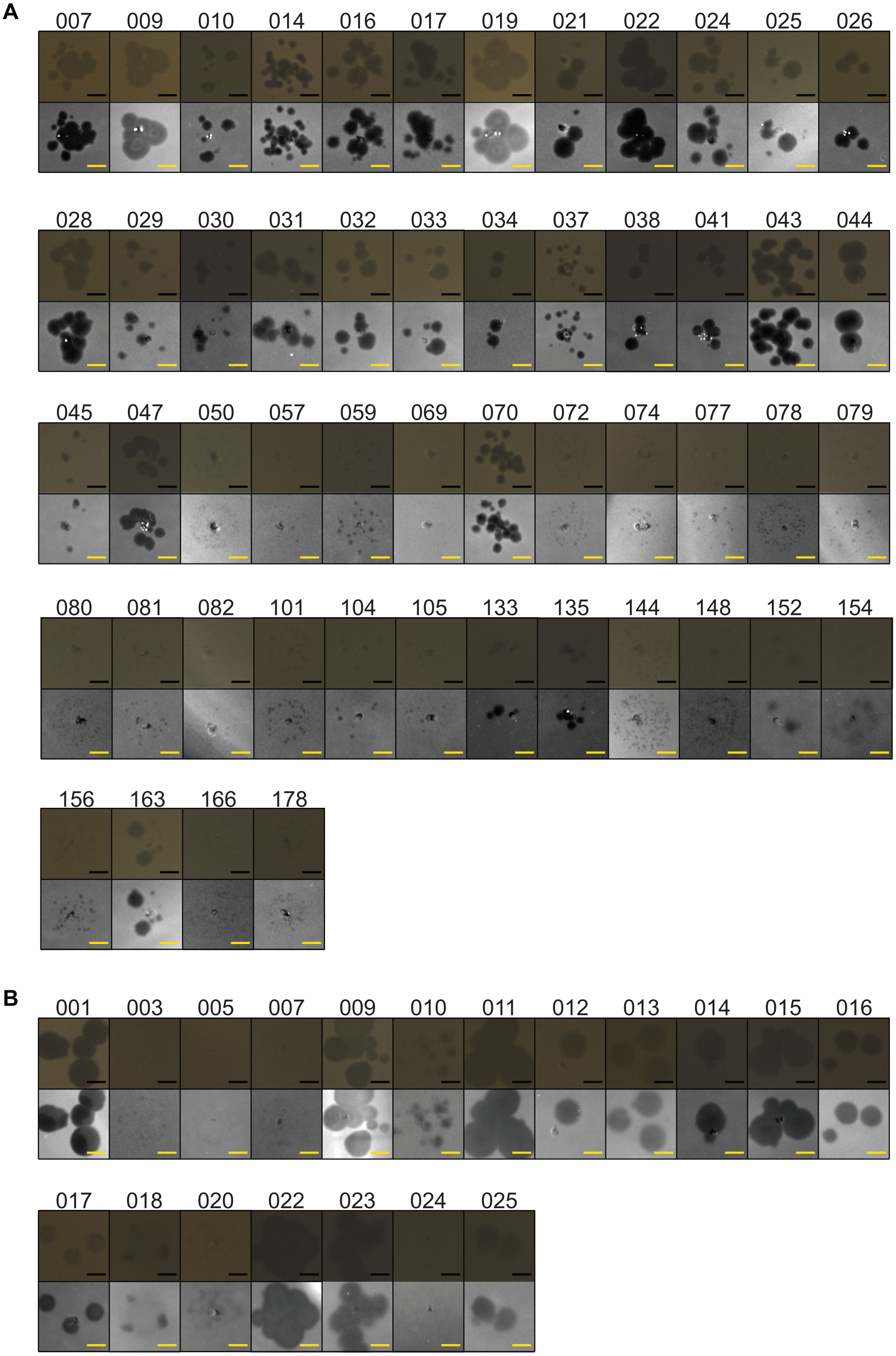

### Supplemental Figure 3

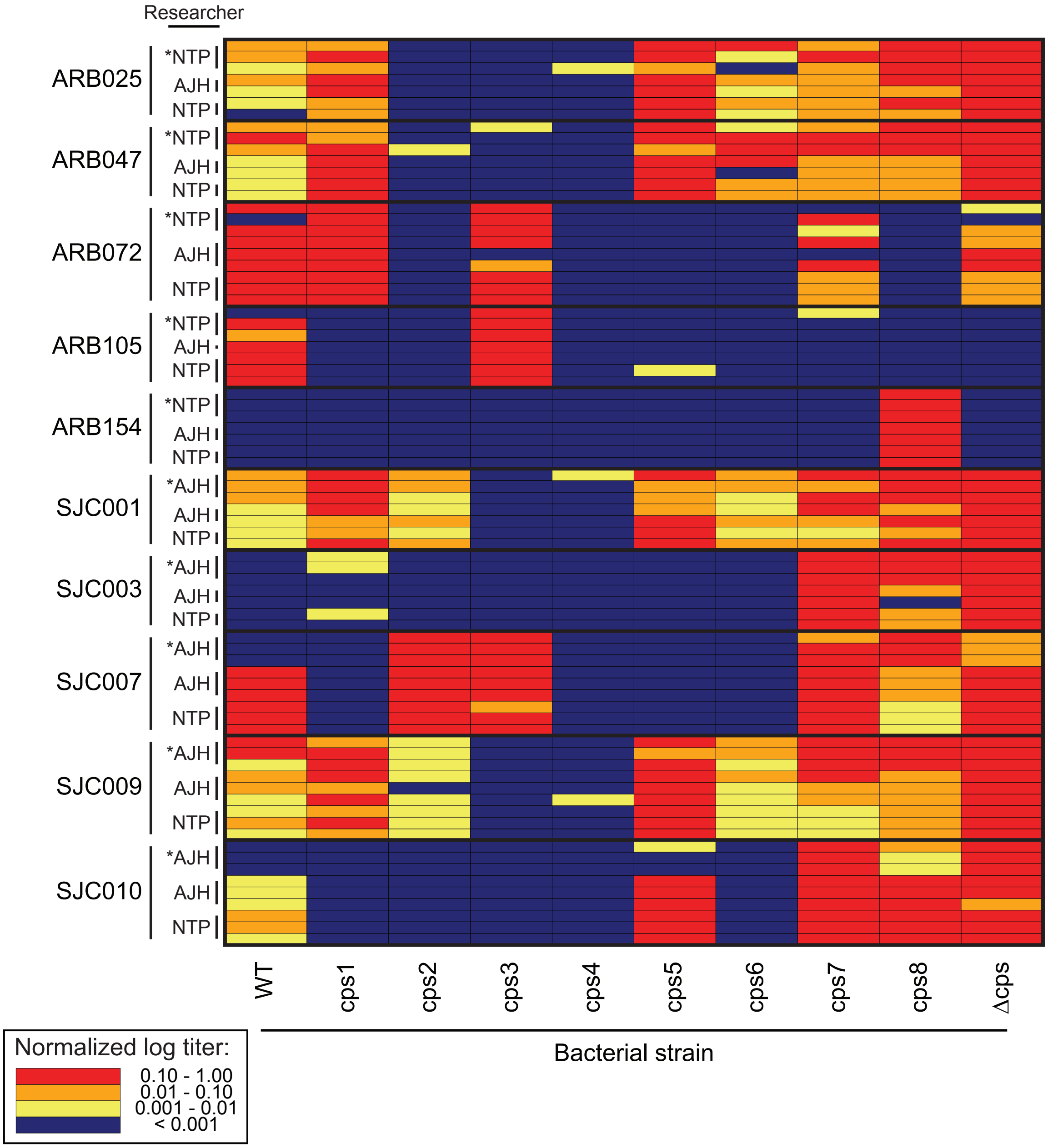

### Supplemental Figure 4

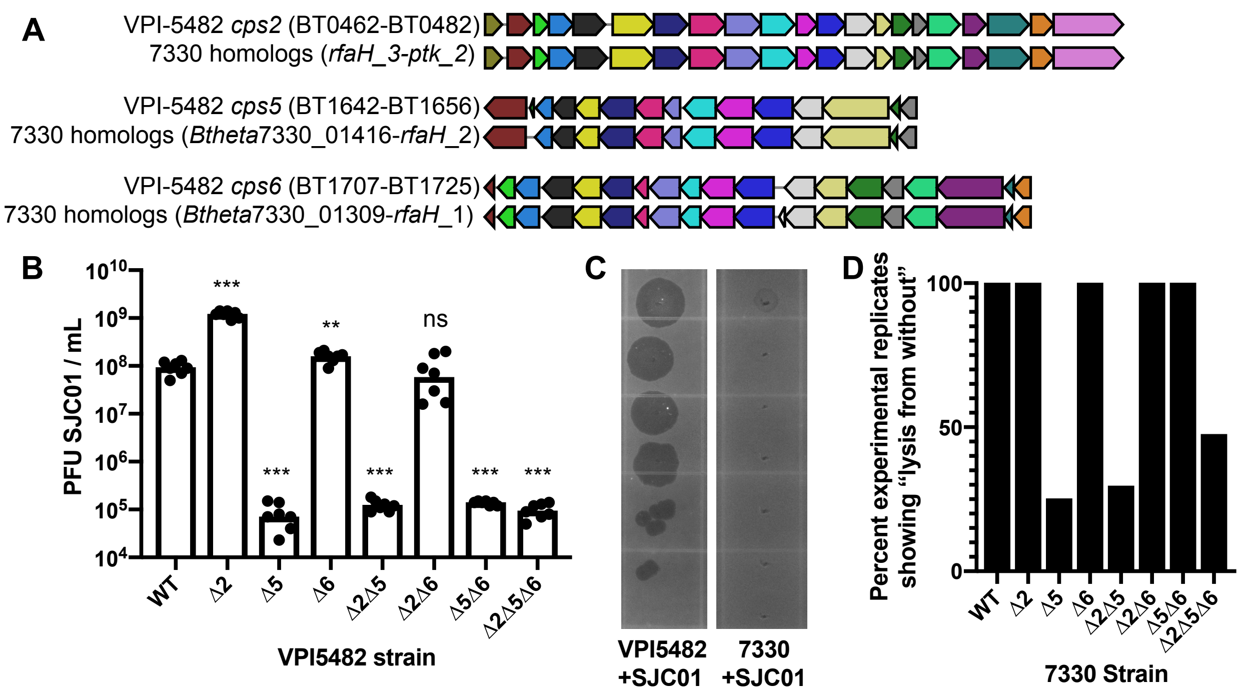

### Supplemental Figure 5

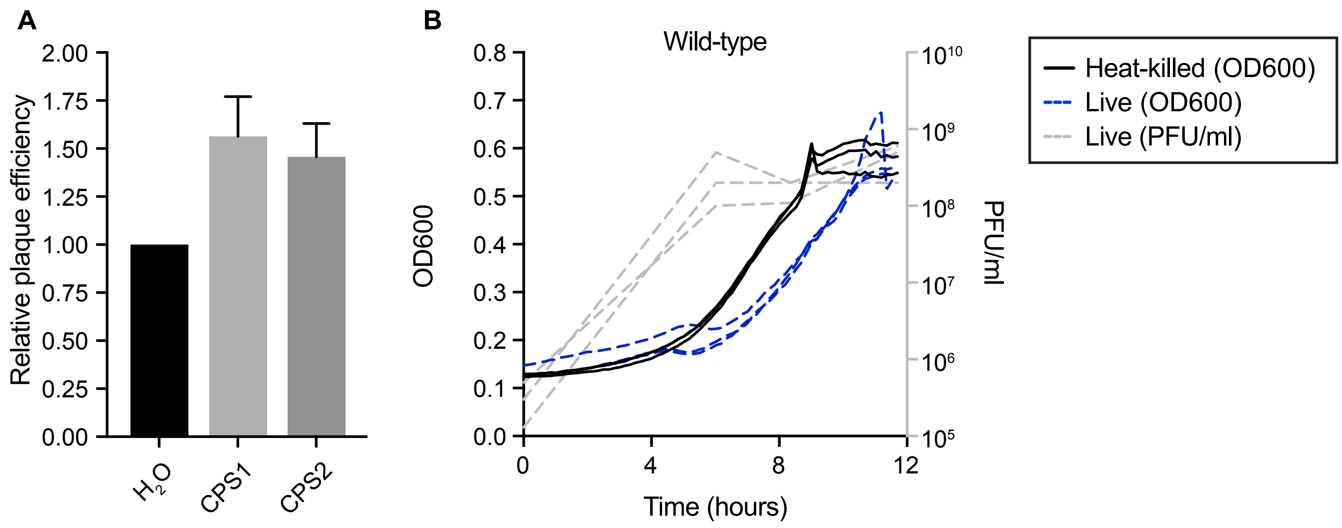

### Supplemental Figure 6

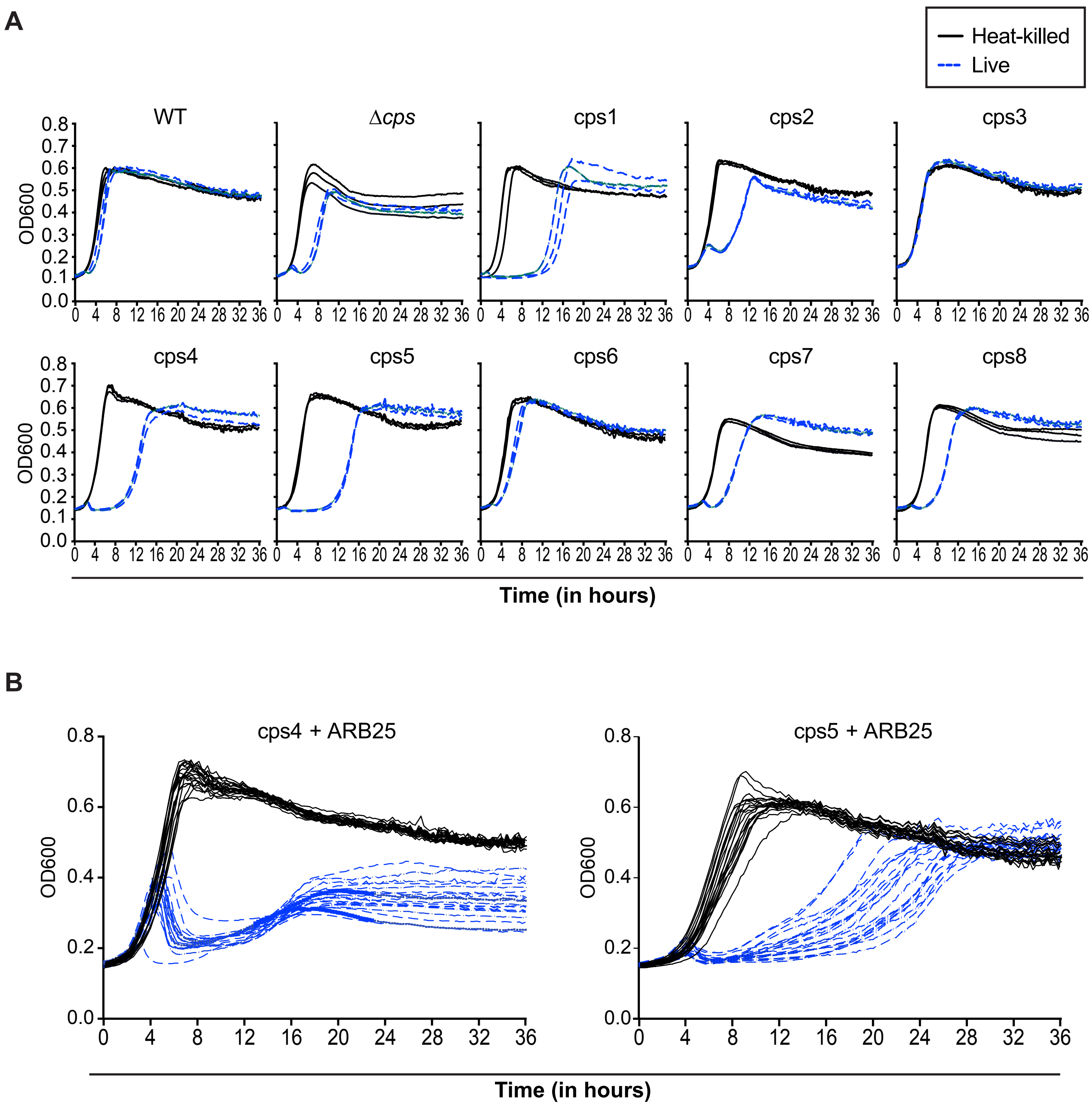

### Supplemental Figure 7

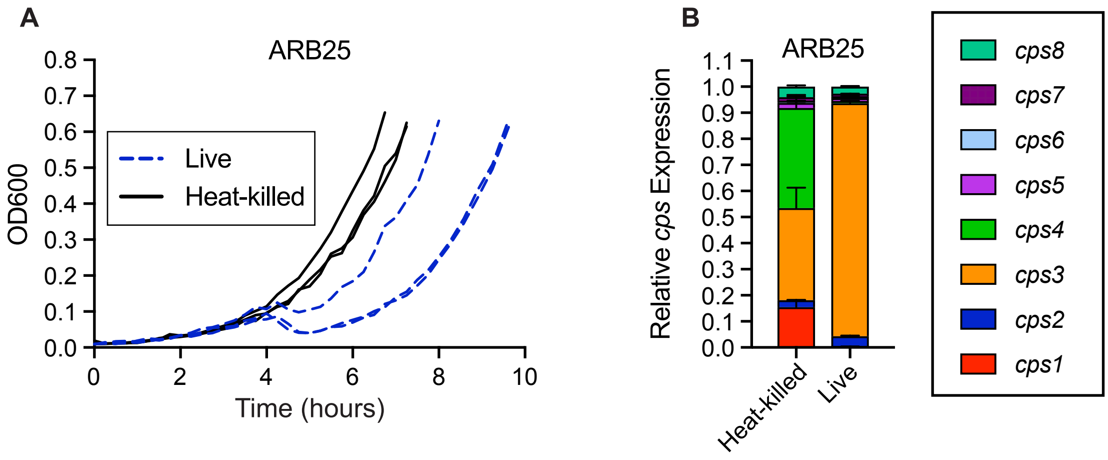

### Supplemental Figure 8

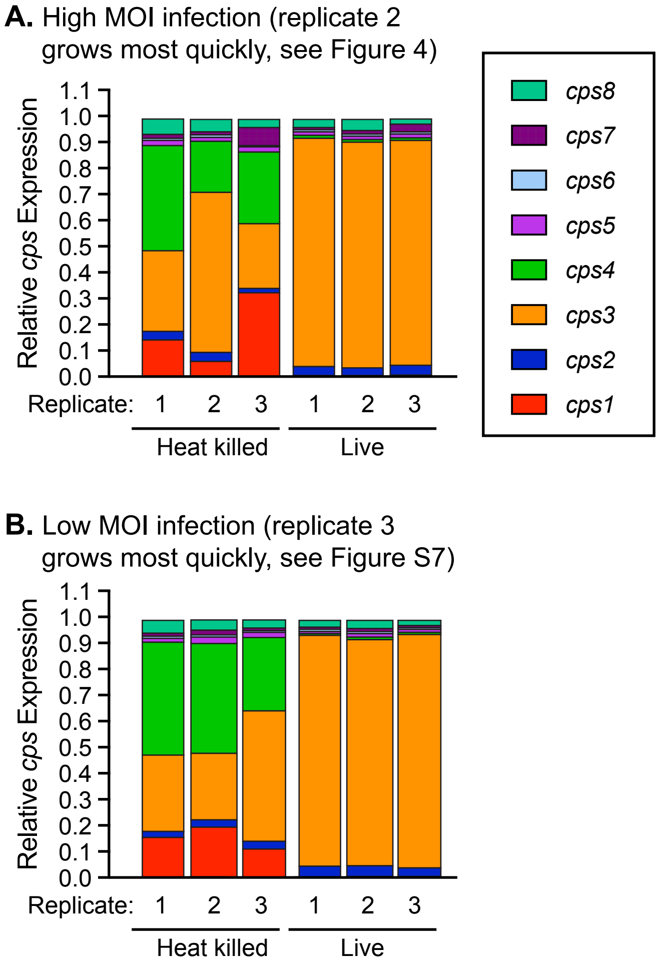

### Supplemental Figure 10

**Figure S10**

A

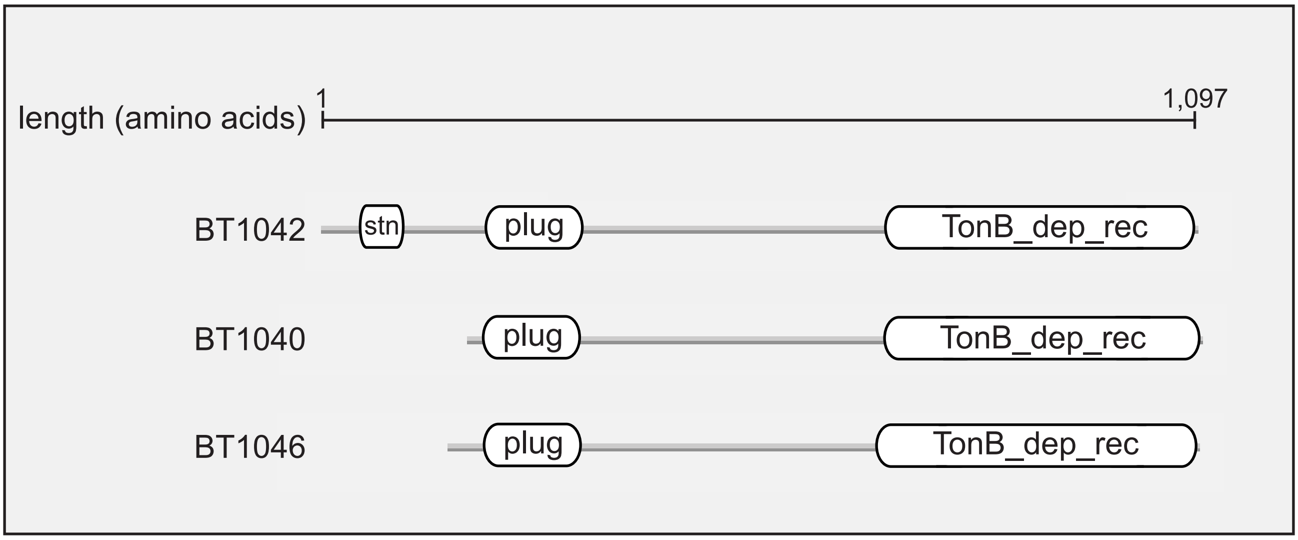


B

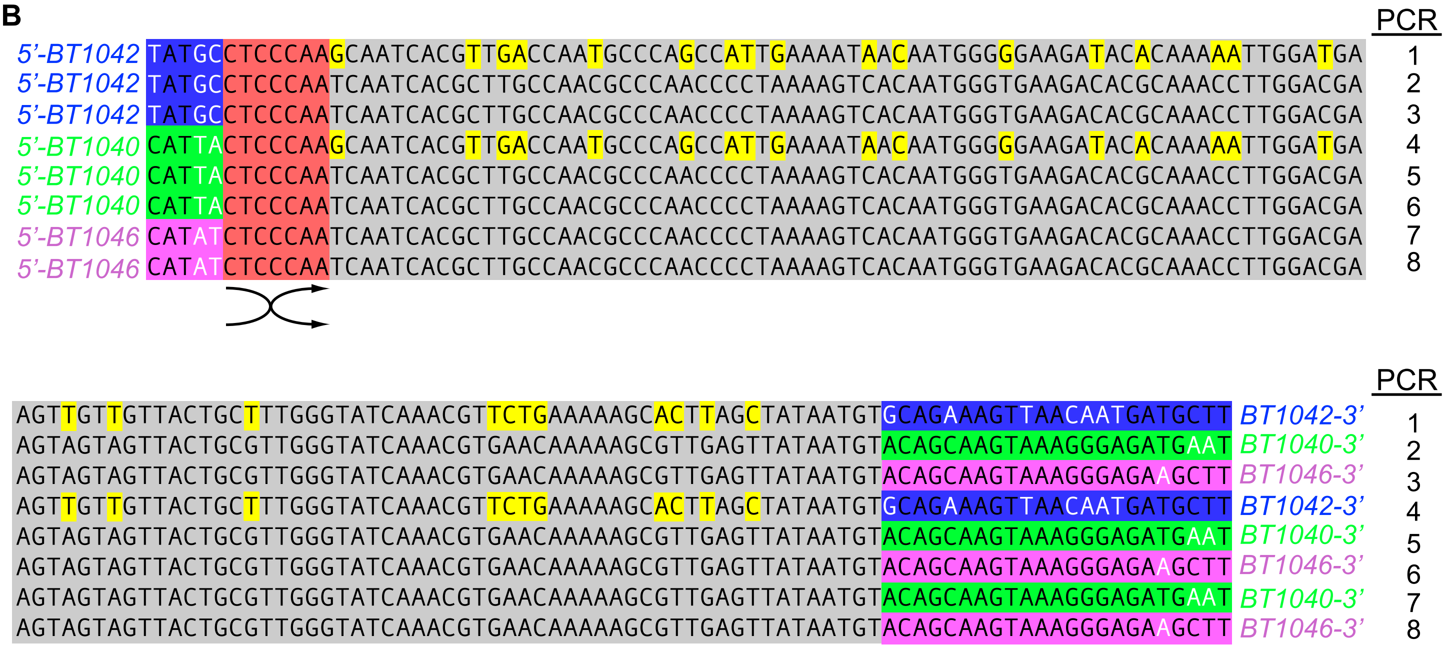

### Supplemental Figure 11

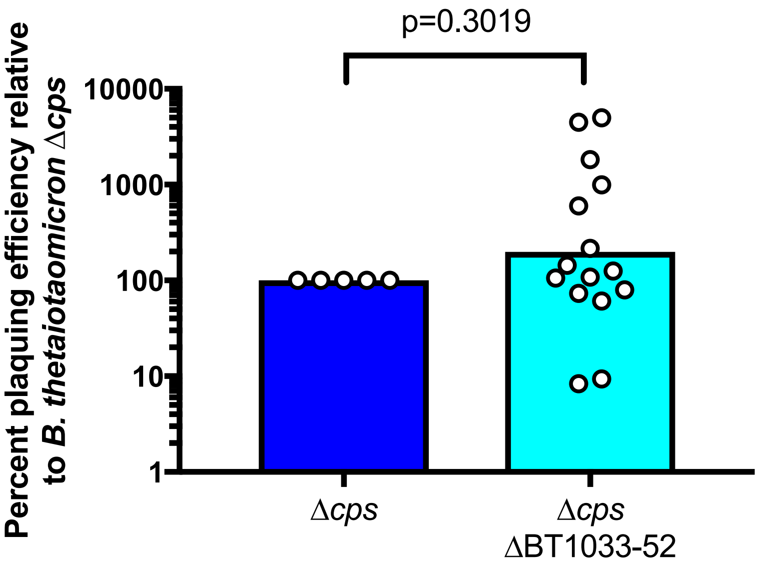

### Supplemental Figure 12

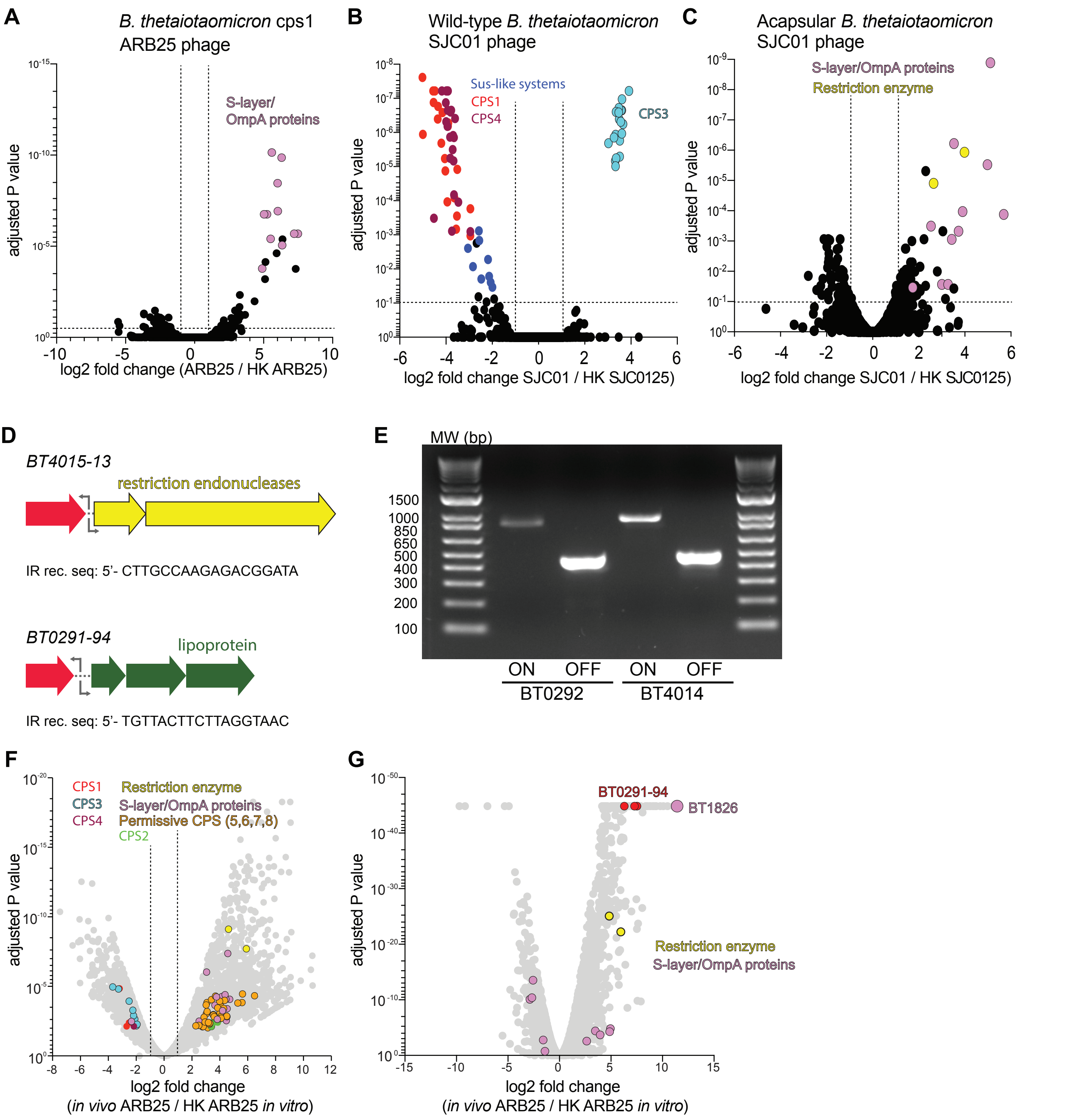

### Supplemental Figure 13

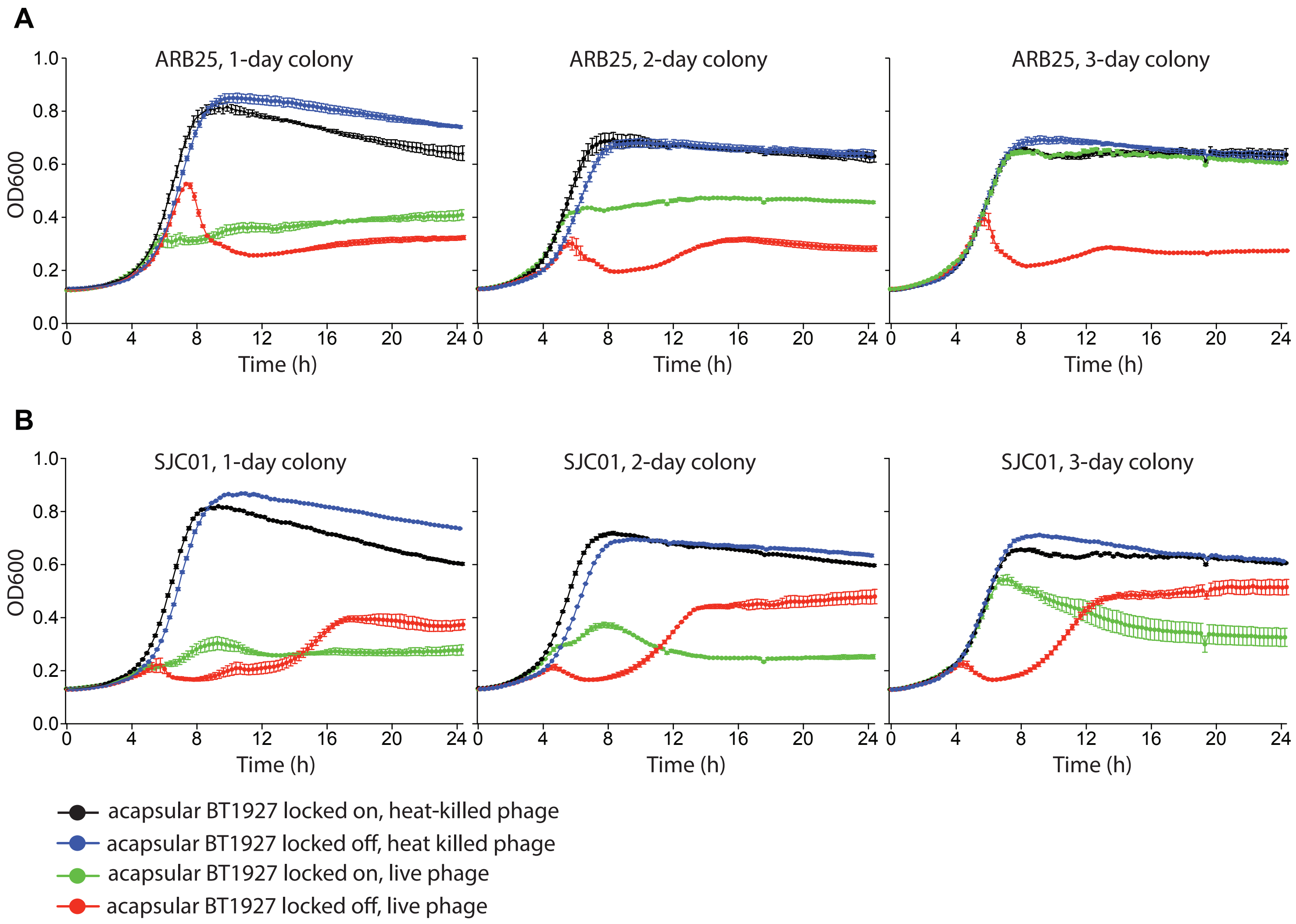
