## Supplemental Figure 9 for "Multiple phase-variable mechanisms, including capsular polysaccharides, modify bacteriophage susceptibility in *Bacteroides thetaiotaomicron*"

**Figure S9**

BT1927 promoter "on" recombination PCR sequence:

new DNA junction (published)

universal primer

recombination site

TTTTGTTTGCATGTTTTCCAAAATATACCCATAtTTGCaGCGTCTTTTCTTCCcGTTaCCTACAAAGTAACGGAA
AGACAGACCACCAAGATGAGATGGATTATGAAATGACAGCAAACAAATTTTTTATTTATTAATTGTATAACATTTAATTT
ACTTTTATGAGAAAATGGACGTATTTGGTCGCAACCTTGCTAATGGCGGGTACGACAGCTACATTCACGGGATGAA

BT1826 promoter "on" recombination PCR sequence:

new DNA junction

universal primer

recombination site

TTTTTGTTTGCATGTTTTCCAAAATATGTCCATATTTGCAGTGCTATTCAGTCCCGTTACCTACCAAGTAACGGGACAGGCATTATTG
AAACTTAGTGTTTAATAATATATTATTAATGTAATTTTTAAGTAGTTATGAAAAGAAAATTTGTAAAAGTGATGTTCTTCGGGGCGTTAG
CACTTTCTACCGTCACCTACGTAGGCTGTAAGGACTACGATGATGACATCAAAAACGTGCAGGAACAGATTGATGCCATCAACAAG
AAAGGTGCTGATGTTACAACTGAAGCAATGACTGCTGCTATTAATAGCGCAGTAGCCGGATTGCAGACACAATTGGATGCTATCGC

BT1502 promoter "on" recombination PCR sequence:

new DNA junction

universal primer

recombination site

TTTTTGgTTGCATGTTTTCCAAAATATCTCCATCTTTGCAACGCTGTTTAGCTCCGTTACCTATGAAGTAACGGGCAAAAAGTATTG
GGAGTAAGTTTCTTGTTCATCCTCTATGGAGCATGGACACAGGTAAAACAAAAAAAGAAAAGGAGGTTACACTAAAAATGAATGGG
GACAACACATGGGATTATTGAATCGACATCGAGAGATTATATTTTTATTATTTAATTTATAAATTAAAAGTTTTTTTATTATGAACAAAAA
GTTTTTAAATGCAGTCCTGTTTGGAGCGTTATTAGCGTCTTCGACGGGCACATTTACATCTTGTAAAGACTACGATGACGACATCA
ATGGTCTGAGCGAACGTGTGGATGCAGTCGAAAAAACATTAGCTGACTTGAATACCAAGTTCGGAGCATTGGCTTATGTGAAATC

BT1507 promoter "on" recombination PCR sequence:

new DNA junction

universal primer

recombination site

TTGTTTGCATGTTTTCCAAAATATATCCATCTTTGCAaCGCTATTCaGCTCCGTTaCCTACAAAGTAACGGGTAAAAAGTATTGGGA
GTAAGTTTCTTGTTCATCCCCCTGTGGAGTATGAACACAAGTAAAACAAAAAAGAAAAGGAGGTTATACTGAAAATGAATGGGGACA
ACACATGAGAGTTTTAAATCAACAATGCGAGACTATATTTTTATTATTTAATTTATAAATTAAAAGTTTTTTTATTATGAACAAAAAGTTTTT
AAGTGTAGTCGTGTTTGGGGCGCTGTTGGCGTCTTCTGCAGGTACATTTACGTCTTGTAAAGACTATGACGACGACATCSATGCAG
TAAACGGCAGAATCGATGAACTCGCAAAAAGTCTTTCAGATTTGCAGGCCAAAGTTGGGTCTTTTGTAAAGTCTGTTACTTATGATC
CGGCTACGGGAAAATTGACTGTGGTAGATGGAGAGAACAAC

BT4481 promoter "on" recombination PCR sequence:

new DNA junction

universal primer

recombination site

CATGTTTTCCAAAATATATCCATCTTTGCAACGCTaTTTAGCTCCGTTACCTAGaAAGTAACGGGTAAAGGCTGTT
TGGCCCGTATGCTTCATAGTGTATTATATAAGGGAAATAAAGGTTGAATGAATTAAGAAAAGGAGGTCGCACGAGAAATGAACGG
AACAACACATGGGATTTAAAATCAAAAGTGAGAAATTATATTTTTATTATTTTTTTAATTTATAAATTAAAAGTTTTTTTATTATGAACAA
AAAGTTTTTAAATGCTGTCCTGTTTGGAGCGCTGATAGCGTCCTCGGCAGGTACATTTACTTCTTGTAAAGACTACGATGMCG
ACATCAAGAGTCTGCAGGAACAAATTGACAAAAGCGGAAGCACCGTTGGCGATCTTCAGACTCAGCTGACAACGCTGAAGA
GTGCTGCAGAAGCTGCAtCAGGCTGCTGCTGATGCTGCAAAGAgCAGCTGCTGCAGAAGCTAAGACTGCCGCAGAAGCTGC

BT2486 promoter "on" recombination PCR sequence:

new DNA junction

recombination site

TCCTTGCGCGTTTTCTcCCCGTT-CCTACAA-GTAACGGGCGAATATTATTTG
AACTTAGTGTTTAATAATATATTATTAATGTAATTTTTAAGTAGTTATGAAGAAAAAATTTATTAGAGTGATGTTTTTCGGAGCATTG
GCGCTAACCACATTTTCATATGTAGGTTGTAAGGACTACGATGACGATATCGATCGGTTAGAGCAGAAGATAACCGAAAATGC
CAACGCTATTGCGGAAATCAACAAGCTGATCGAAGGGGGATCTGTGATCACAAAAGTTGAGAACATTACCGGTGGTGTGGA
AGTAACATTGAGCAACGGTAAGAGCTTTACCATTAAAAATGGTGATGCCGGAAAAGATGGTACTCAGTGGACGATTGGTGAA
GACGGATACTGGTATCAAGATGGTAAAAA
